# Qombucha: Reconstructing unobserved progenitor methylation profiles reveals distinct developmental programs in glioblastoma

**DOI:** 10.64898/2026.08.06.743400

**Authors:** Xuan Cindy Li, H. Lalchungnunga, Ananth Hari, Yuelin Liu, Omkar Singh, Zhichao Wu, Zied Abdullaev, Stephen M. Mount, Kenneth D. Aldape, Eytan Ruppin, Alejandro A. Schäffer, S. Cenk Sahinalp

## Abstract

Glioblastoma (GBM) is a highly aggressive brain cancer characterized by substantial intratumoral heterogeneity. Previous research demonstrates that GBM may have complex cell origins. To elucidate the interplay between brain development and GBM progression, we introduce **Qombucha** (Quadratic prOgraMming Based tUmor deConvolution with cell HierArchy), a computational framework that uses DNA methylation data to infer tumor cell-type composition and profiles of unobserved progenitor cells. Unprecedentedly, **Qombucha** incorporates a developmental cell hierarchy that models mature brain cell types and their progenitors.

Applied to a large TCGA GBM dataset spanning the RTK I, RTK II, and MES TYP subtypes, **Qombucha** identifies a distinct cell type composition profile for each subtype and recapitulates known biological patterns, including elevated microglia infiltration in MES TYP tumors and increased RTK I stemness. It also identifies subtype-specific developmental programs and shows that higher progenitor-cell abundance is associated with poorer survival. **Qombucha**-imputed cell fractions map methylation profiles of tumor samples to a compact, 11-dimensional latent space; in an independent NCI GBM cohort, this compact representation improves subtype clustering and enables accurate subtype classification, achieving performance comparable to stateof-the-art models based on full methylation profiles with much higher dimensionality. These results suggest that **Qombucha**-imputed tumor cellular composition captures the core biological axes along which GBM subtypes diverge.

## INTRODUCTION

Glioblastoma (GBM), one of the most aggressive and common malignant brain tumors, has poor prognosis due to its invasive nature and resistance to conventional therapies (Hubert and Lathia 2021; Ostrom et al. 2020). Treatment failures in GBM are attributed to the limitations of doing brain surgery (Cote et al. 2017) and to high heterogeneity both within and across GBM tumors (Sottoriva et al. 2013; Patel et al. 2014; Richards et al. 2021). The intertumor heterogeneity of GBM can be characterized by differences in molecular markers, including DNA, mRNA, and methylation (Zhang et al. 2020). Verhaak et al. (2010) pioneered mRNA-based classification of GBM and established the Classical, Mesenchymal, Proneural, and Neural subtypes. These transcriptomic subtypes exhibit different responses to therapies (Verhaak et al. 2010), although genomic analyses showed that multiple subtypes are often present in the same patient Sottoriva et al. (2013); Neftel et al. (2019). To an extent, prognosis can also be predicted via the presence/absence of somatic alterations in genes such as *IDH1*, *IDH2*, *PTEN*, and *EGFR* (Houllier et al. 2010; Zhang et al. 2020).

There has been a growing focus on DNA methylation-based GBM classification and subtyping (Kumar et al. 2018; Zhang et al. 2020; Koelsche and von Deimling 2022). Sturm et al. (2012) characterized six DNA methylation-based GBM subtypes: K27, G34, IDH1, RTK (receptor tyrosine kinase) I, RTK II, and MES (mesenchymal), among which RTK I, RTK II, and MES are most frequent in adult GBM (Drexler et al. 2023). These three DNA methylation-defined subtypes have been associated with differences in incidence of seizures, spatial location within the brain, genomic features, and tumor purity. RTK I tumors are mainly found in the right frontal lobe of the brain (Foltyn-Dumitru et al. 2024) and are enriched for platelet-derived growth factor receptor-alpha (*PDGFRA*) gene amplifications. RTK II tumors are often found in the left-hemispheric regions (Foltyn-Dumitru et al. 2024) and are associated with higher seizure incidence (Ricklefs et al. 2022) and frequent epidermal growth factor receptor (*EGFR*) gene amplifications. MES tumors are commonly observed in the left-hemispheric regions (Foltyn-Dumitru et al. 2024), tend to have lower tumor purity and neurofibromin-1 (*NF1*) aberrations and increased level of immune cell infiltration (Chen and Hambardzumyan 2018). Using a subset of 32k methylation probes with high variance across subtypes, Capper et al. (2018) extended this line of work to obtain a classifier for 98 brain and central nervous system (CNS) tumor subtypes, including GBM subtypes RTK I, RTK II, and MES.

Single-cell RNA-seq data have improved genomic understanding of GBM Patel et al. (2014); Venteicher et al. (2017); Weng et al. (2019); Neftel et al. (2019); Couturier et al. (2020), but all the larger datasets rely on bulk measurements. In this study, we compile from TCGA and analyze a dataset of 896 IDH-wildtype (no somatic mutation in *IDH1* or *IDH2*) GBM tumors with bulk methylation data. We also obtained an independent IDH-wildtype GBM cohort from the National Cancer Institute comprising 556 patients. For comparison between IDH-wildtype and IDH-mutant tumors, we also compile from TCGA a third dataset of 959 IDH-mutant tumors.

For analyzing bulk methylation data from a large cohort, we introduce a new method, **Qombucha**, to address: how can one estimate the proportion of each cell type from among a defined set (next paragraph and Methods) of cell types and their unseen progenitors? Cell type estimation problems/solutions are generally called “deconvolution”. For bulk transcriptomics data, there are various deconvolution methods some of which jointly estimate the proportion of each cell type and the proportion of total expression of a gene attributable to that cell type Newman et al. (2015); Becht et al. (2016); Aran et al. (2017); Finotello et al. (2019); Newman et al. (2019); Wang et al. (2022); Zaitsev et al. (2022).

The deconvolution of bulk methylation data has also been studied by others using the following equation, where *B* is the observed bulk data.

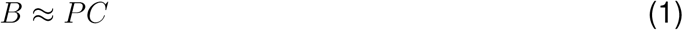

The prior work on methylation data deconvolution falls into three categories: (i) supervised (reference-based), (ii) unsupervised (reference-free), and (iii) semi-supervised (hybrid) (Ferro dos Santos et al. 2024). The supervised methods rely on the known methylation profiles of cells of interest, represented by the known matrix *C*, to estimate the cell type compositions, represented by the unknown matrix *P* in Equation (1). Houseman et al. (2012) conducted pioneering work in methylation deconvolution and developed a linear constrained projection-based algorithm that estimates cell type proportions in bulk sample data by minimizing the differences between the observed bulk sample data and the projection of a reference matrix onto the cell-type proportion matrix. Other methods, such as MethAtlas (Moss et al. 2018) and MethylResolver (Arneson et al. 2020), have employed least squares regression. In addition, machine learning approaches have been developed, including the support vector regression-based MethylCIBERSORT (Chakravarthy et al. 2018) and the deep neural network-based cfSort (Li et al. 2023). The unsupervised methods, in contrast, have been developed for situations where both *P* and *C* are completely unknown and need to be inferred simultaneously. The unsupervised methods are typically based on non-negative matrix factorization (NMF), including MeDeCom (Lutsik et al. 2017) and EDec (Onuchic et al. 2016), or on expectation–maximization (EM) algorithms, such as MethylPurify (Zheng et al. 2014) and PRISM (Lee et al. 2019). The semi-supervised methods combine unsupervised techniques and prior knowledge. In these approaches, *C* includes full reference panels for known, pre-specified cell types, but also leaves room for additional cell types lacking a reference panel. CelFiE (Caggiano et al. 2021), for example, uses a block coordinate descent-like algorithm to infer the compositions for known cell types and a user-defined number of unknown cell types. PREDE (Qin et al. 2020) and PRMeth (He et al. 2022), on the other hand, are iterative NMF-based deconvolution methods that jointly estimate cell-type proportions and the profiles of unknown cell types (expression for PREDE; methylation for PRMeth). Both adopt an all-or-nothing assumption about reference profiles: for each cell type, the profile is either fixed (known) or entirely free (unconstrained), which can lead to overfitting.

Methylation deconvolution has been applied in GBM to elucidate the tumor’s epigenetic landscape. Drexler et al. (2024) used a reference-free deconvolution method developed by Silverbush et al. (2022) to elucidate cell types and cell states in bulk GBM methylation data. A follow-up paper by Silverbush et al. (2026) used a similar reference-free deconvolution method to identify three distinct malignant cell states and concluded that RTK I cases are enriched with (the more stem-like) cell state 1. Wenger et al. (2019) used the R package InfiniumPurify (Zheng et al. 2014) to deconvolve GBM tumor methylation data and assess tumor purity. Another study by Iluz et al. (2026) compiled the methylation profiles of differentiated non-neuronal cell types from previous studies, and used them together with an experimentally determined bulk methylation profile of *in vitro*-induced oligodendrocyte precursors to identify the prevalence of each such cell type in GBM samples. However, none of these prior studies specifically addressed the question of deconvolving GBM methylation data to explore GBM subtype-specific brain cell development processes. Importantly, the previous applications of methylation-based deconvolution generally considered observable differentiated cell types but not the unobservable progenitors.

In contrast, our analysis of cell types in GBM is focused on the six *non-neuronal differentiated cell types* studied by Tian et al. (2023) as well as their *progenitors*. We additionally included the excitatory and inhibitory neuronal cell types as they likely introduce impurities in GBM samples (Auld and Robitaille 2003). The six differentiated non-neuronal cell types of most interest include: astrocytes (ASC), endothelial cells (EC), microglia (MGC), oligodendrocytes (ODC), oligodendrocyte precursors (OPC), and (the union of) vascular and leptomeningeal cells (VLMC). Studies of various tissue types and organs support the notion that developmental programs, which differentiate unobservable progenitor cells to observable mature cells are branched processes (Patel and Yanai 2024; Gong et al. 2025; Domcke and Shendure 2023). Therefore, we consider the observable and progenitor cell types as a tree that can aid in the deconvolution because methylation status is typically inherited from parent cell (type) to child cell (type). Considering various cell types simultaneously is justified by previous research showing that GBM has complex cell origins, including neural stem cells (NSC) and non-neuronal cells like ASC and OPC (Zong et al. 2015). GBM tumors also display diverse cell states with transcriptomic signatures that are ASC-like, neural progenitor cell(NPC)-like, OPC-like, and MES-like (Neftel et al. 2019).

After we estimated cell type proportions, we investigated associations between cell types and tumor types using the methylation-based classification method of Capper et al. Capper et al. (2018), IDH mutation status, and overall survival. We also investigated which methylation sites give the most signal for inferring cell type proportions and whether the methylation status of those sites is stable in cell differentiation, as explained below.

Previously, Tian et al. (2023) performed single-cell whole genome bisulfite sequencing (scWG seq) for 40 major brain cell types in 46 brain regions across samples from three healthy males and reported that 800 features grouped from 156k CpG sites are sufficient to differentiate mature brain cell types whose developmental program was represented as a tree. It is possible to trace the origin of each cell by identifying those CpG sites that go through methylation status change in distinct stages of development/differentiation Gabutt et al. (2022). Therefore, we used methylation features in Tian et al. (2023) to define our tree-based deconvolution problem below. Because the individuals from whom the samples were used for the model in Tian et al. (2023) are healthy, we avoid the potential circularity of using GBM-based modeling in our approach.

Analysis of a normal development tree process in cancer data could be confounded by the established theory of Nowell (1976) that subclones within solid tumors such as GBM also evolve in a tree-like manner. The study of inferring the tree history of tumors from static snapshots of tumor omics data is called “tumor phylogenetics”; most tumor phylogenetics studies to date have been based on somatic mutations or copy-number variants Malikic et al. (2015); Beerenwinkel et al. (2015); Schwartz and Schäffer (2017); Li et al. (2024). Some tumor phylogenetics studies have instead used methylation data as the input and found that evolution of “tumor lineage informative” CpG sites also forms a tree Bian et al. (2018); Cresswell et al. (2020); Liu et al. (2023); Gabbutt et al. (2025). The overlay of a methylation-based tree structure for normal development and another methylation-based tree structure for tumor progression could be problematic if the same (un)methylated CpG sites are most informative for both trees. We show early in *RESULTS* that the methylation site overlap is not substantial in a statistical sense.

Because the methylation profiles of progenitor cell types in the developmental tree are not available Tian et al. (2023), **Qombucha** employs the empirical observation that, for many features, a progenitor’s methylation state matches that of its mature descendants when those descendants agree. We show that the tree inheritance property enables the deconvolution of the bulk GBM methylation data by jointly estimating each sample’s cell-type composition and the unknown progenitor methylation values across the cohort. Intuitively, this allows one to attribute portions of the bulk signal to specific developmental paths (cell-of-origin/trajectory) while preserving tumor-evolutionary information carried by lineage-informative CpG sites Bian et al. (2018); Capper et al. (2018); Irizarry et al. (2009); Liu et al. (2023); Gabbutt et al. (2025); Tian et al. (2023).

In *RESULTS*, we formalize the problem that **Qombucha** solves and summarizes our key results in the application of **Qombucha** to a GBM dataset we compiled from TCGA, including those pointing to distinct cell type compositions of the GBM subtypes. We also present results from IDH-MUT glioma patient data compiled from TCGA and a novel GBM data set generated at the National Cancer Institute (NCI). For the NCI GBM dataset, we assigned a GBM subtype to each tumor sample using a classifier trained on the 11 cell type fractions that **Qombucha** inferred from the TCGA GBM dataset. Notably, this classifier matched the accuracy of a state-of-the-art classifier based on the 10,000 most variable methylation sites. We further included results from simulated data for evaluating the robustness and efficiency of **Qombucha**. The *DISCUSSION* summarizes the main contributions of this paper and lays out a plan for future work. Section *METHODS* gives a detailed description of **Qombucha**, including the key algorithms, preprocessing of inputs, design of our simulations, and other methods used in our biological and computational analyses.

## RESULTS

### Summary of Main Findings

We applied **Qombucha** to bulk methylation data from 896 GBM patients from TCGA spanning multiple subtypes and an additional 959 IDH-MUT patients and showed that **Qombucha** successfully recapitulated the established biological insight that GBM MES TYP patients exhibit higher immune cell infiltration (Chen and Hambardzumyan 2018) and that GBM RTK I tumors exhibit an increased level of stemness (Silverbush et al. 2026; Matsumoto et al. 2025; Wang et al. 2021). We further validated our findings on an independent GBM cohort of 556 patients collected and sequenced at NCI. To validate or analysis of the TCGA data set, we began by applying the TCGA-derived **Qombucha** reference panel to deconvolve the NCI samples. We then trained a GBM subtyping classifier using the 11 **Qombucha**-inferred cell-type fractions from the TCGA cohort and assigned subtypes to the NCI GBM samples based on their inferred cell-type compositions. Remarkably, the classifier matched the performance of a state-of-the-art approach based on the 10,000 most variable methylation sites.

Beyond recovering known biology and demonstrating cross-cohort generalizability, **Qombucha** identified GBM subtype-specific brain developmental programs and demonstrated that these programs are significantly associated with patient outcomes. Importantly, the **Qombucha-** imputed mature and progenitor cell fractions provide a set of low-dimensional and interpretable features yielding subtype clusters that are “as tight” or “tighter—than” those clusters derived from high-variance methylation features commonly used in clinical classifiers Capper et al. (2018). Our work offers a framework for understanding subtype-specific GBM progression at a granular level and provides insights that may help refine future precision medicine for GBM.

### Brief Description of Qombucha

The input to the key computational problem we aim to solve is a matrix *B* consisting of bulk methylation feature values per patient, and a matrix *C_p_* of partially known brain cell methylation profiles: *C_p_* includes the fully known methylation profiles of mature cell types (Tian et al. 2023), and “copies” to a progenitor the methylation feature value of its descendant mature cells, only if these cells agree in that feature value (Section *Cell hierarchy-informed partial inference of progenitor cell type profiles*) otherwise the feature value in the progenitor remains unknown. *B* has dimensions *n × m*; each row represents a patient and each column represents a methylation feature defined by Tian et al. (2023), which is a set of CpG sites similarly methylated across distinct cell types. Our goal is to factor *B* approximately into: (i) a cell composition matrix *P* of dimensions *n × r*, where each column is a cell type and each row consists of inferred fractions of cell types per patient sample, and (ii) a complete reference panel matrix *C_f_*, of dimensions *r × m*, where each row contains methylation feature values for each mature and progenitor brain cell type under consideration, with the constraint that the known entries of *C_p_* are preserved in *C_f_* . Because *C_f_*represents the completed version of the partially known input matrix *C_p_*, we represent either matrix with *C* when there is no ambiguity. The subscript *p* stands for “partial” and the subscript *f* stands for “full” or “filled in”.

The matrix factorization problem to simultaneously infer brain progenitor profiles and cell composition in bulk methylation data, can be formulated as a Quadratic Program (QP). This formulation leverages the known brain cell type hierarchy to inform the establishment and partial inference of progenitor cell profiles. The resulting partially complete reference panel *C_p_*, together with the bulk methylation data matrix *B*, is the input we use to deduce the unknown entries in *C_p_* and the cell type composition matrix *P* .

The objective of the QP is to minimize the matrix reconstruction error, specifically, the sum of the squared differences between the input bulk methylation matrix *B* and *P × C_f_* . The rows of matrix *C_p_* pertaining to observable brain cell types are known, while the rows corresponding to the progenitor cells can only be partially characterized from the leaves of the tree *T* denoting the brain developmental program.

Due to the computationally intensive nature of simultaneous inference of the two matrices, and the existing limitations of commonly used optimization tools such as Gurobi, the implementation to directly solve this QP is not yet scalable. Thus, we developed **Qombucha**, a block coordinate descent (Wright 2015)-based approach that initializes the unknown entries in the reference panel *C_p_* by assigning them those values that minimize the differences between the unknown entries of the progenitor cell types- and those of their descendant cell types, as indicated by the brain development tree *T* . The resulting matrix *C*_0_, kickstarts an iterative process to infer matrices *P* and *C_f_* . In each iteration *ι* = 1, 2*, . . .*, **Qombucha** first uses matrix *C_ι_*_−1_, to infer the cell composition matrix *P_ι_*using the QP formulation. Next, **Qombucha** uses *P_ι_*to infer *C_ι_*, again by using the QP formulation, to estimate the full reference panel *C_f_*, until convergence. Because only one of the matrices needs to be inferred in each step of the iteration, our QP formulation works efficiently.

Figure 1 shows the key deconvolution problem of our study, and Figure 2 demonstrates the overall workflow of **Qombucha** to solve this problem. A detailed description of **Qombucha**, the input preparation process (including, e.g., adding methylation profiles of other cell types to handle sample impurity), as well as the algorithm to obtain *C*_0_ are provided in *METHODS*. An open source implementation, which requires a Gurobi (www.gurobi.com) license, is available at https://github.com/algo-cancer/Qombucha.

**Figure 1:**
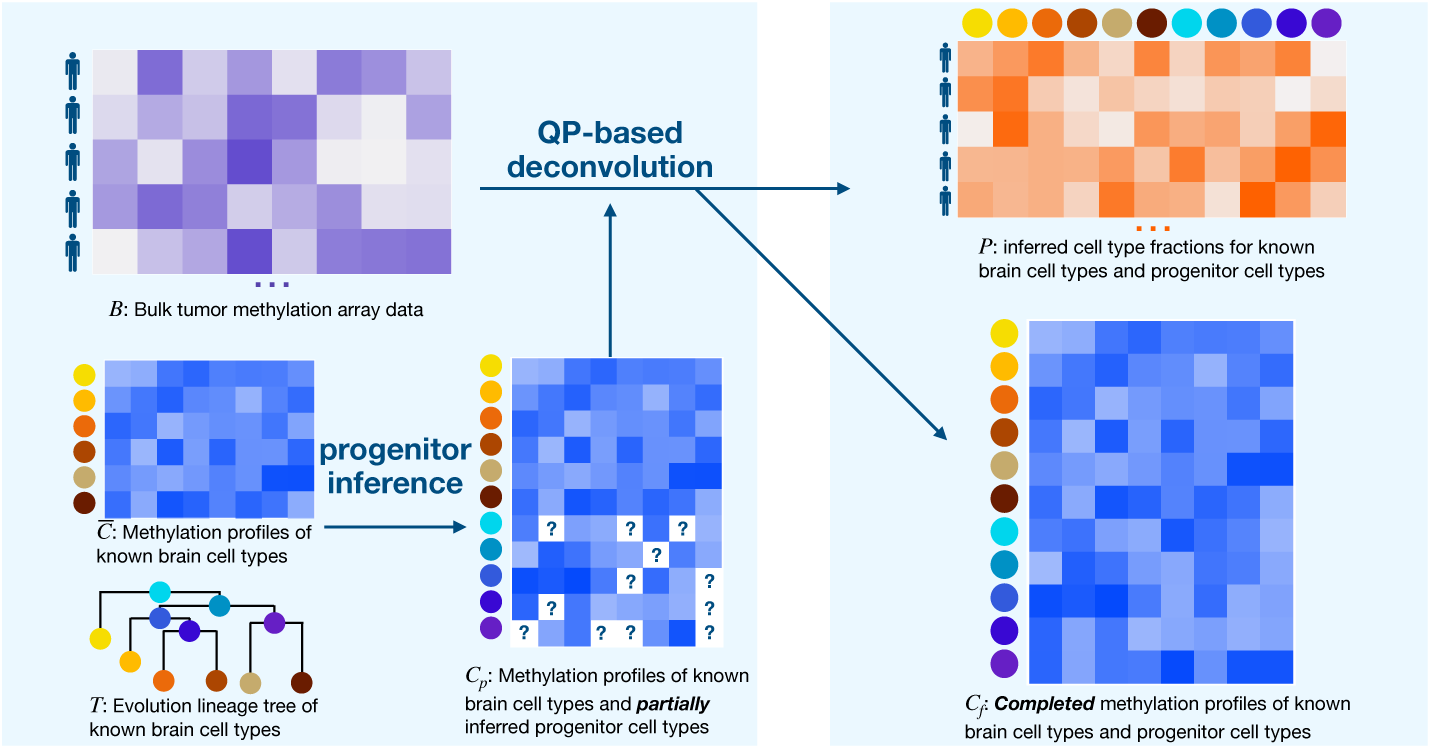
The key computation problem solved by Qombucha formulated as Quadratic Programming (QP). The input to the QP consists of: (i) a partially complete reference panel *C_p_* which includes the complete methylation profiles for known developed brain cell types and partially complete profiles for progenitors, and (ii) the bulk methylation data matrix *B*. The QP asks to infer the full reference panel *C_f_* and the cell type composition matrix *P* with the objective of minimizing the sum of the squared differences between *B* and *P × C_f_* .

**Figure 2:**
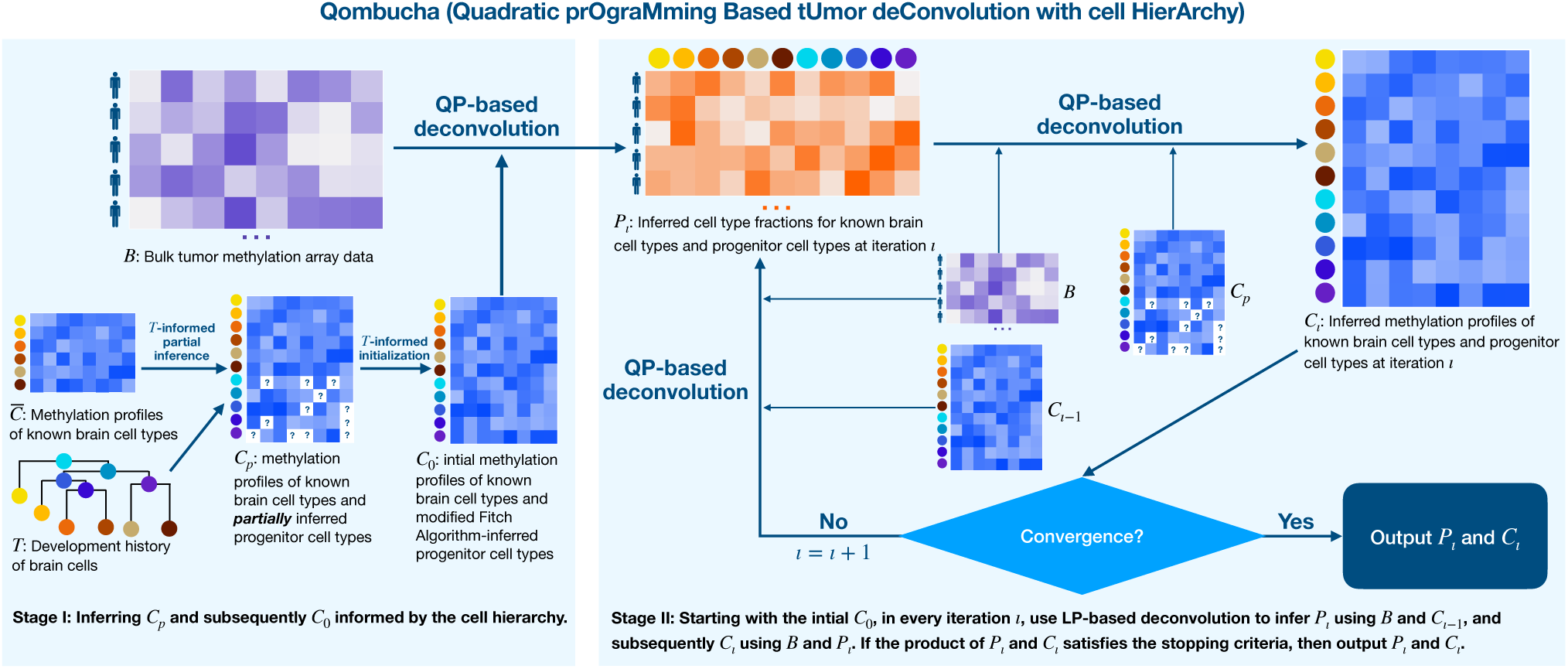
Graphical abstract of Qombucha’s block coordinate descent-like approach. Qombucha. first builds the initial reference panel *C*_0_ by imputing the unknown entries of *C_p_*—i.e., the methylation features of the progenitor cell types for which the descendant mature cell methylation values do not agree—to minimize progenitor–descendant differences under the developmental tree *T* . Then, for iterations *ι* = 1, 2*, . . .*, **Qombucha** alternates QP subproblems: infer cell proportions *P_ι_* from *C_ι−_*_1_, then update the unknown entries of *C_p_*to obtain *C_ι_* using *P_ι_*. Upon meeting the stopping criterion, **Qombucha** outputs *P* = *P_ι_* and *C_f_* = *C_ι_*. Because *P_ι_* is inferred from fully determined *C_ι−_*_1_ and *C_ι_* is subsequently inferred from the fully determined *P_ι_*, the QP steps are more efficient than the QP to infer *P* and *C_f_*simultaneously.

### Methylation Sites Associated with Brain Cell Development and Brain/ CNS Cancer Subtyping

Our **Qombucha**-based analysis for GBM overlays a methylation-based tree structure for normal development and that for abnormal tumor progression. Because this overlay would be problematic if the two tree structures relied on many of the same methylations sites, we quantitatively evaluated whether the CpG sites that are associated with brain cell development/differentiation are mostly distinct from those that are associated with specific brain/CNS cancer subtypes.

We first show that the genomic regions of the brain development/differentiation associated methylation sites identified by Tian et al. (2023) and those of the brain/CNS cancer associated methylation sites identified by Capper et al. (2018) are distinct: the development-associated sites are less concentrated in CpG islands and shores and more concentrated in inter-CGI than the brain-cancer subtype associated sites (Figure 3A).

**Figure 3:**
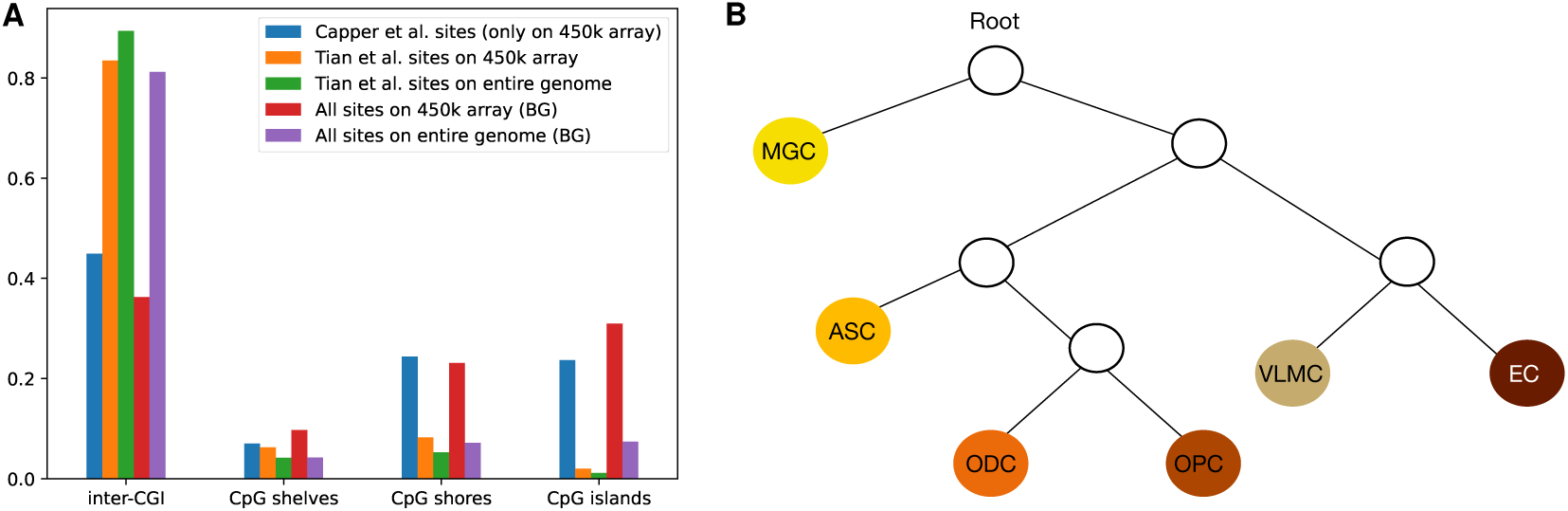
Methylation sites associated with brain cell development/differentiation differ from sites that characterize brain/CNS cancer subtypes. **(A)** Comparison of the genomic regions of brain cell type-differentiating sites and brain cancer subtype-differentiating sites. Four categories of genomic regions are present in the bar plot: CpG islands (CGI), CpG shores (2kbp spanning CGI), CpG shelves (2kbp spanning the shores), and inter-CGI regions. The blue bars indicate the 32k CpG sites differentiating brain cancer subtypes identified in Capper et al. (2018) in each genomic region category. These sites are all on the Illumina 450k methylation array because Capper et al. (2018) only included array data in their analysis. The green bars indicate the 156k CpG sites differentiating normal brain cell types reported in Tian et al. (2023). The orange bars indicate a subset of the 156k CpG sites overlapping with the Illumina 450k methylation array. The red bars indicate all CpG sites on the Illumina 450k methylation array, serving as background (BG) fractions. The purple bars indicate all CpG sites in the entire human genome, also serving as background fractions. **(B)** Brain development history represented as a tree. The methylation sites associated with brain development, which is distinct from those associated with brain cancer subtypes, suggest a tree structure for representing the brain development history, inferred by the method in Section *Reconstructing brain development history using single-cell methylation data*. The colored leaf nodes represent the developed brain cell types whose methylation profiles are fully known. The internal nodes (with no labels) represent the progenitor cells whose methylation profiles are partially derived by **Qombucha** from their descendant brain cell types.

Next, we demonstrate that the brain development/differentiation-associated methylation sites identified by Tian et al. (2023) are statistically distinct from the brain/CNS cancer subtype associated sites used by Capper et al. (2018). Of the 485,512 sites in the 450k methylation array, 32,000 have been reported by Capper et al. (2018) to be associated with brain/CNS cancers. Among the 156,000 sites identified by Tian et al. (2023) as mature brain cell type-differentiating, 2484 overlap with the 450k methylation array. Among these 2484 sites, only 108 overlap with Capper et al.’s 32,000 sites; in constrast, a random selection of 2484 sites from the 450k array would have an expected overlap of 32000 *×* 2484*/*485512 = 163.72 with the Capper et al. (2018) sites. The p-value of observing an overlap of *≤* 108 sites between a set of 2484 sites and another set of 32, 000 sites is Pr(*X ≤* 108) = 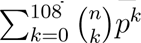 (1 *− p*)*^n^*^−*k*^ = 1.136 *×* 10^−6^, where *X* is the binomial random variable representing the number of overlaps, 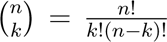 is the binomial coefficientfor *k* successes in *n* trials, and *p* = 32000*/*485512 *≈* 0.06588 is the probability that a randomly selected 450k methylation array site is among the Capper et al. (2018) 32,000 sites.

The above observations suggest that the methylation sites that mark brain cell differentiation are substantially distinct from those for brain/CNS tumor progression.

### Biological and Clinical Insights

In this section, we present our key biological and clinical findings derived from **Qombucha**- imputed cell fractions and progenitor methylation profiles. Our methodological contribution is to utilize the prior knowledge of brain cell developmental history to guide the inference of progenitor cell methylation profiles, which, to the best of our knowledge, has not been employed by previous attempts at methylation deconvolution. Our work addresses a different question compared to prior deconvolution efforts. We aim to identify whether distinct GBM subtypes are associated with distinct mature or progenitor cell types, as well as distinct developmental trajectories, i.e., specific paths in the tree representing the developmental program of brain cells.

To evaluate the generalizability of **Qombucha**-derived biological insights, we used the TCGA GBM cohort to infer a complete cell-type reference panel and applied it to deconvolve bulk methylation profiles from an independent NCI GBM cohort. We then used the TCGA-derived **Qombucha** cell-type fractions to train a GBM subtype classifier and assigned subtypes to the NCI tumor samples based on their inferred cell-type compositions.

### Qombucha recapitulates known GBM subtype characteristics

We first obtained the brain development history (Section *Reconstructing brain development history using single-cell methylation data*, Figure 3B) and subsequently reference panels *C_p_* and *C*_0_ (Section *Obtaining the partially complete reference panel C_p_ and initial reference panel C*_0_ *for non-neuronal cell types*). Using *C*_0_, *C_p_*, and *B*, the bulk methylation array data of TCGA 896 GBM patient samples (processed as described in Section *Processing bulk sample methylation data*) as input, we employed **Qombucha** to obtain deconvolved cell type compositions per sample. The completed reference panel *C_ι_* obtained at convergence was then used to perform *one-step* deconvolution of the bulk methylation data of the NCI cohort.

As can be observed in the third column of Figure 4A and Figure 4B, in both the TCGA and NCI cohorts, compared to RTK I and RTK II, the patient samples in the MES TYP subtype demonstrate a higher presentation of microglia, which are the resident immune cells in the brain (Soulet and Rivest 2008). This result recapitulates the known association with higher immune activity and inflammation in the tumor microenvironment and increased levels of immune cell infiltration in the MES TYP subtype (Chen and Hambardzumyan 2018).

**Figure 4:**
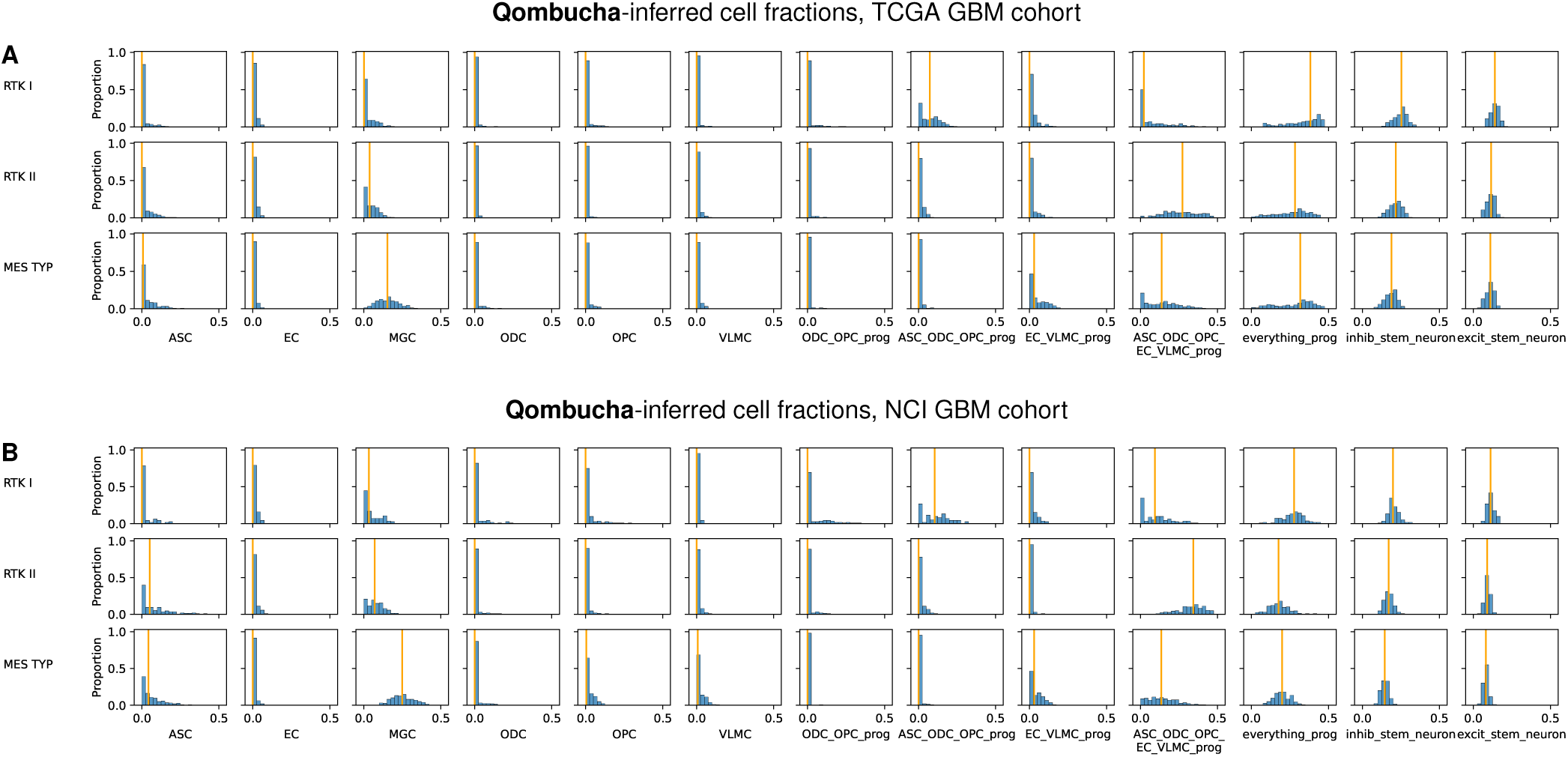
Comparing cell proportions imputed from the TCGA cohort and those from the NCI cohort. Each row represents a specific GBM subtype: RTK I, RTK II, or MES TYP, and each column represents a specific cell type. The orange vertical line in each plot indicates the median inferred fraction (across all samples for the associated GBM subtype represented by that row) of the cell type represented by that column. **(A)** Cell type compositions inferred by **Qombucha** in each GBM subtype in the TCGA cohort. **(B)** Cell type compositions inferred by **Qombucha** in each GBM subtype in the NCI cohort. See Figure S4 for the Kolmogorov-Smirnov (KS) tests comparing each GBM subtype against the remaining two subtypes on **Qombucha**-inferred cell fractions in each patient cohort.

We further evaluated the stemness of each GBM subtype using the estimated proportion of the “everything progenitor” as a proxy. By this measure, in both TCGA GBM and NCI GBM cohorts, RTK I has a significantly higher stemness than either MES TYP (TCGA GBM: *P <* 9e-11, NCI GBM: *P <* 5e-19, Wilcoxon test) and RTK II (TCGA GBM: *P <* 9e-21, NCI GBM: *P <* 5e-21); MES TYP has more stemness than RTK II (TCGA GBM: *P <* 0.013. NCI GBM: *P <* 0.00021). This order of stemness among the three subtypes is consistent with two recent methylation-based analyses of Matsumoto et al. (2025) and Silverbush et al. (2026). An earlier expression-based analysis of stemness by Wang et al. (2021) ranked the subtypes as RTK I *>* RTK II *>* MES TYP in decreasing order of stemness. These results show that **Qombucha** is able to confirm previously discovered biological insights on GBM subtypes.

### Qombucha reveals that brain developmental programs are similarly GBM subtype-specific in both cohorts

We show in Figure 4 that different brain developmental programs, represented by cell type fractions, appear to be associated with different GBM subtypes in both the TCGA cohort and the NCI cohort. To assess whether the differences in imputed cell type fractions across GBM subtypes are significant, for each cohort, we performed two-sided Kolmogorov-Smirnov (KS) tests (Kolmogorov 1933; Simard and L’Ecuyer 2011). Specifically, we compared the fractions imputed for patients of one GBM subtype against the imputed fractions of patients of the other two subtypes. The p-values obtained from these tests in Figure S4 show that MGC is the most significantly different cell type across all GBM subtypes in both cohorts and distinguishes the MES TYP subtype from other subtypes, reinforcing our observations described in Section ***Qom- bucha*** *recapitulates known GBM subtype characteristics*.

We also evaluated whether the imputed cell fractions of the NCI cohort have similar distributions compared to those of the TCGA cohort. As can be seen in Figure 4, the distribution of cell proportions in each GBM subtype in the NCI cohort (Figure 4B) is highly similar to that of the TCGA cohort (Figure 4A). To quantitatively evaluate this similarity, we performed Wilcoxon rank-sum tests on the ranking of the inferred cell proportions (see Figure S5 and Section *Wilcoxon rank-sum test for evaluating similarity of cell proportions across cohorts*). All but one of the 39 comparison has a p-value larger than the Bonferroni-correted false discovery threshold, demonstrating that **Qombucha**-imputed cell proportions are robust in different patient cohorts.

### Qombucha-inferred cell fractions enable accurate classification of GBM subtypes

We evaluated the ability of **Qombucha**-inferred cell type fractions to classify GBM subtypes using the ensemble framework described in Section *Classification of GBM subtypes using **Qombucha**-inferred cell fractions*. Specifically, we obtained the complete cell type reference panel *C_ι_* from the TCGA GBM cohort and used *C_ι_* to perform one-step deconvolution on the independent NCI GBM cohort of 556 patients to assess generalizability. The reference subtype annotations (“ground truth”) for both the TCGA and NCI cohorts were based on the DNA methylation classifier developed by Capper et al. (2018), which assigns CNS tumor subtypes based on the 32,000 most variable CpG sites. These reference subtype labels served as the benchmark for evaluating Qombucha’s GBM subtype classification performance.

Classification performance on the TCGA cohort was assessed using the 5 *×* 5 nested crossvalidation framework described in the Methods (*Classification of GBM subtypes using **Qombuch** inferred cell fractions*). For external validation, subtype predictions for the independent NCI cohort were generated by averaging the outputs of all 25 trained models, without additional training or parameter optimization.

On the TCGA dataset, the ensemble classifier achieved an overall accuracy of 88% (787/896) and a macro-averaged AUC of 0.96 (Figure 5A,D). When applied to the independent NCI cohort, the model achieved an accuracy of 91% (509/556) and a macro-averaged AUC of 0.98, demonstrating strong transferability across datasets (Figure 5B,E). Receiver operating characteristic (ROC) analysis demonstrated high discriminative performance across all three GBM subtypes. Confusion matrix analysis further confirmed robust classification accuracy across all subtypes. The MES subtype was classified with particularly high accuracy, with a misclassification rate of 2.7% (7/258). The RTK II subtype showed a moderate misclassification rate of 10.2% (19/186), while the RTK I subtype exhibited a higher misclassification rate of 18.8% (21/112) (Figure 5B). Although the Capper et al. (2018) classifier remains the clinical standard for methylation-based CNS tumor classification, a recent study suggested that classifiers trained using 10,000 highly variable CpG sites can provide improved biological resolution and more refined characterization of CNS tumor classification Sill et al. (2026). Therefore, we evaluated a classifier trained directly on the 10,000 most highly variable CpG sites to compare its classification performance with that achieved using Qombucha-inferred cell type fractions. Importantly, the classification performance achieved using **Qombucha**-inferred cell type fractions was comparable to that obtained using high-dimensional (i.e. with 10,000 CpG sites) DNA methylation array data (Figure 5C,F). See Section *Classification of GBM subtypes using highly variable methylation CpGs* for details.

**Figure 5:**
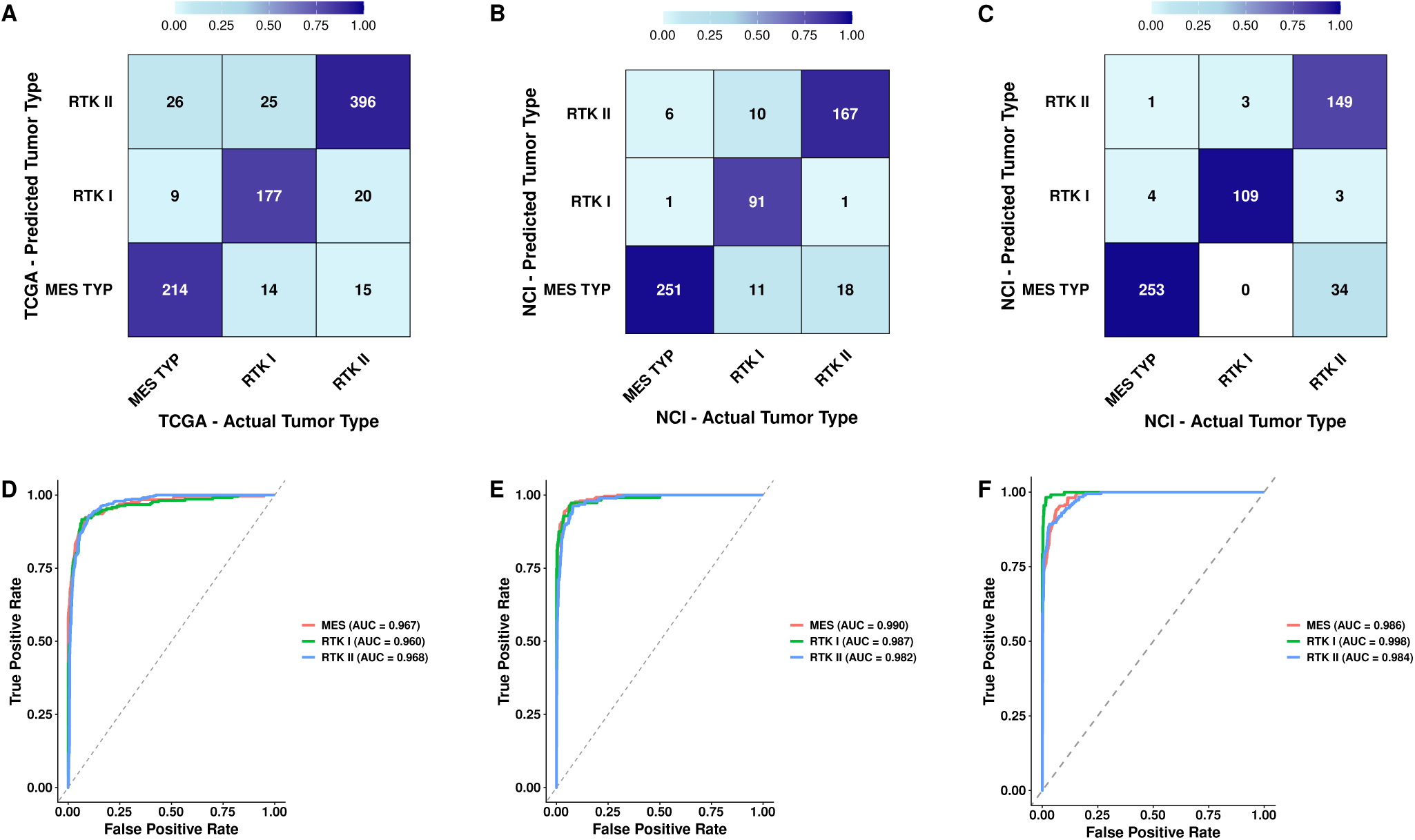
Classification performance of glioblastoma subtypes using Qombucha-inferred cell proportions and DNA methylation profiles. **(A)** Confusion matrix for glioblastoma subtype classification (MES, RTK I, and RTK II) in the TCGA training dataset using **Qombucha**-inferred cell proportions. **(B)** Confusion matrix for glioblastoma subtype classification in the NCI validation dataset using **Qombucha**-inferred cell proportions. **(C)** Confusion matrix for glioblastoma subtype classification in the NCI dataset using the top 10,000 most variable DNA methylation probes. **(D)** ROC curves demonstrating classification performance in the TCGA training dataset (area under the curve (AUC) = 0.965). **(E)** ROC curves demonstrating classification performance in the NCI validation dataset (AUC = 0.986). **(F)** ROC curves demonstrating classification performance using the top 10,000 most variable DNA methylation probes in the NCI dataset (AUC = 0.989).

Despite relying on a substantially lower-dimensional feature space, the **Qombucha**-based classification approach achieves similar accuracy and AUC, highlighting the ability of **Qombucha** to capture the key biological signals necessary for subtype discrimination. Overall, these results demonstrate that **Qombucha**-inferred cell type compositions provide a highly predictive feature representation for GBM subtype classification that generalize to an independent cohort.

### Qombucha-imputed cell type fractions improve GBM subtype clustering compared to state-of-the-art methods

We evaluated the tightness of known GBM subtype clusters in the TCGA GBM cohort using both the *D_in_/D_out_* score and the silhouette score and different choices for the inter-sample distance, as described in Section *Evaluating tightness of the known GBM subtype clusters*. Briefly, both the *D_in_/D_out_*score and the silhouette score *s*(*i*) of sample *i* denote how well the sample is clustered. Smaller *D_in_/D_out_*scores indicate better clustering, whereas larger silhouette scores indicate better clustering. The purpose of this comparison was to assess how a distance based on **Qombucha**-inferred mature and progenitor cell fractions performs in clustering.

We used six distinct types of information to calculate (*L*^1^) distances between all pairs of patients: (i) the methylation values of 2484 (number of Tian et al. (2023) sites on the Illumina 450k methylation array) sites randomly sampled from the 450k methylation array, (ii) the methylation values of the 32k sites used in Capper et al. (2018), (iii) the methylation values of 98 features containing cell type-differentiating sites included in the 450k array (Section *Input data prepara- tion*), (iv) QP-inferred mature brain cell type fractions, excluding progenitor profiles, (v) MethylCIBERSORT(Chakravarthy et al. 2018)-inferred mature brain cell fractions, excluding progenitor profiles, and (vi) **Qombucha**-inferred mature and progenitor cell fractions with a number of distinct starting points. The (*L*^1^) distances are then used to calculate the *D_in_/D_out_* score and the silhouette score *s*(*i*) for each sample *i* for assessing how close the sample is to the other samples are in the same GBM subtype, against those samples in the other two subtypes. Evaluated by both scores, **Qombucha**-inferred mature and progenitor cell fractions for all GBM samples outperform all other types of values in tightness of the known GBM subtype clusters (Figure 6 and Figure S3, row VIII, column “All”).

**Figure 6:**
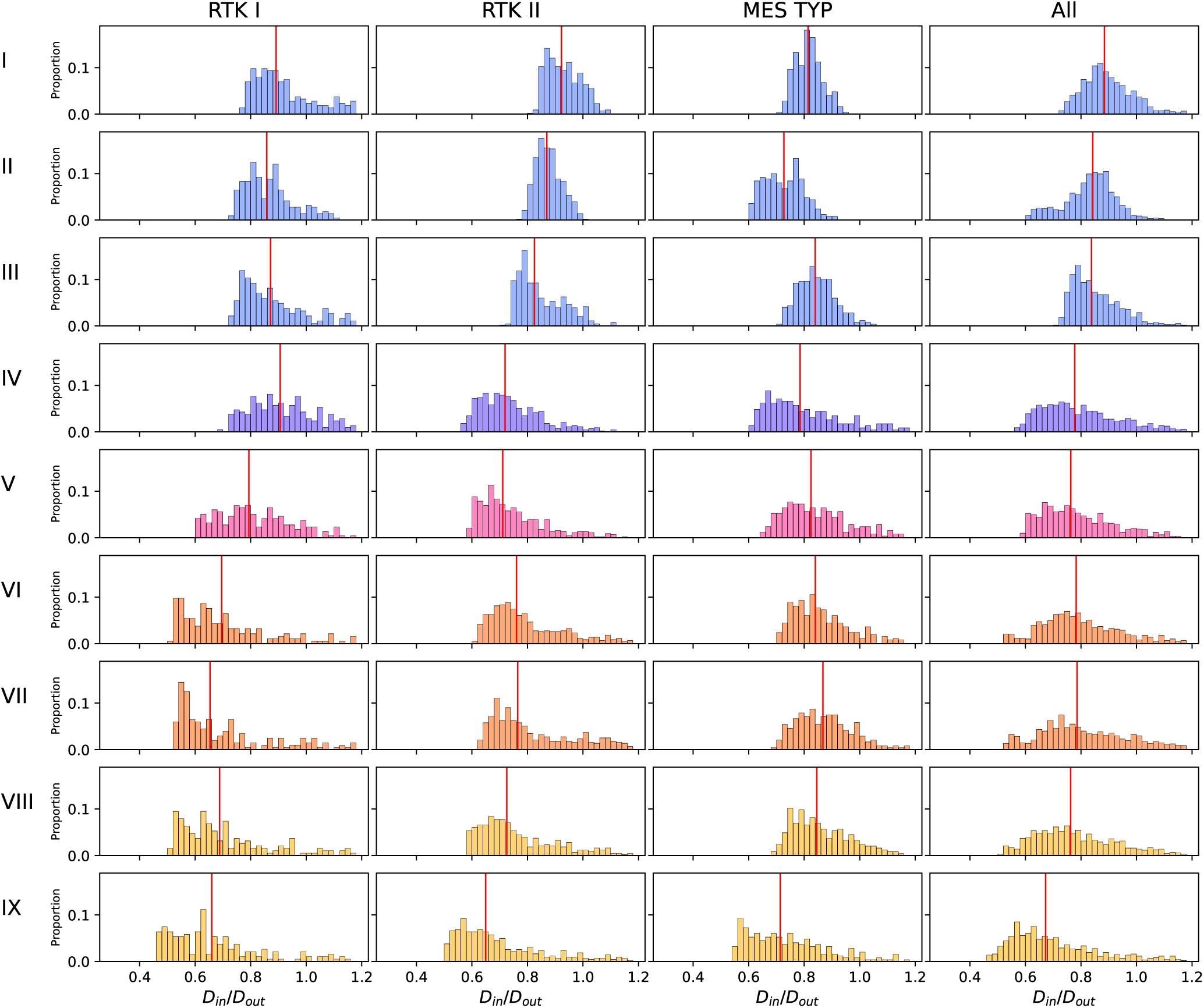
Tightness of GBM subtype clusters computed using. *D_in_/D_out_* **in the TCGA cohort.** Here *D_in_/D_out_*is the ratio between the mean *L*_1_ distance among sample pairs within a specific GBM subtype and that between the samples of this GBM subtype and the remaining samples. Each row uses distinct information to compute the distances for assessing the tightness of each GBM subtype cluster (a smaller *D_in_/D_out_* indicates a better clustering): (I) the methylation values of 2484 sites randomly sampled from 450k methylation array; (II) the methylation values of Capper et al. 32k sites; (III) the methylation values of 98 features; (IV) QP-inferred cell fractions, excluding progenitors; (V) MethylCIBERSORT-inferred cell fractions; (VI) **Qombucha**-inferred cell fractions with progenitor profiles initialized as the mean of direct descendants; (VII) **Qombucha**-inferred cell fractions with random initialization; (VIII) **Qombucha**-inferred cell fractions with development tree-informed starting point; (IX) **Qombucha**-inferred cell fractions with development tree-informed starting point, score calculated from the union of cell type quadruplets best distinguishing each subtype. The scores of IV-VIII are calculated using all 11 cell types including mature nonneuronal cell type and their progenitors as listed in the brain development tree, excluding the impurities consisting of two neuronal cell types. The red vertical lines indicate the median for each subtype in each experiment.

These results show that **Qombucha**-imputed cell fractions of mature and progenitor cells (excluding impurities consisting of the two neuronal cell types), despite being only 11-dimensional vectors, achieve better clustering of GBM subtypes than higher-dimensional methods, such as the Capper et al. (2018)-selected 32k sites (experiment VIII vs. II). Moreover, the **Qombucha**imputed cell fractions outperform GBM subtype clustering based on the original bulk methylation data *B* (experiment VIII vs. III). Conceptually, **Qombucha** embeds each sample in an 11- dimensional space in a manner that is useful for downstream analysis tasks such as clustering and tumor classification. This is surprising since the CpG sites that **Qombucha** considers are associated with the developmental program and have minimal overlap with the CpG sites used by Capper et al. (2018) for brain tumor classification. We observed that it is essential to include progenitor cell profiles in the reference panel *C*, as clustering performance greatly improves when progenitor profiles are included (experiment VIII vs. IV).

Furthermore, we identified a subset of cell types that best distinguish GBM subtypes. We enumerated all possible quadruplets of cell types and calculated the *D_in_/D_out_*using the **Qombucha**-imputed fractions of cell types in each quadruplet. The quadruplets best distinguishing each GBM subtype (i.e., quadruplets with the lowest *D_in_/D_out_*scores for each subtype) are the following: RTK I is best distinguished by MGC, ODC, VLMC, and ASC-ODC-OPC-EC-VLMC progenitor; RTK II by ODC, OPC, ODC-OPC progenitor, and ASC-ODC-OPC progenitor; MES TYP by EC, VLMC, ODC-OPC progenitor, and ASC-ODC-OPC progenitor. Combining the quadruplets that best distinguished each GBM subtypes results in 8 cell types: EC, MGC, ODC, OPC, VLMC, ODC-OPC progenitor, ASC-ODC-OPC progenitor, as well as ASC-ODC-OPC-EC-VLMC progenitor. We calculated the *D_in_/D_out_*using these 8 cell types and demonstrate that the **Qombucha**-inferred fractions of 8 cell types have the best performance in improving the clustering tightness of the GBM subtypes (Figure 6, row IX).

### Qombucha recapitulates known GBM IDH-WT and IDH-mut glioma differences and cell-of-origins of IDH-mut glioma

One of the most important advances in GBM genomics in the 21st century was the recognition that some tumors have recurrent somatic mutations in either of the paralogous genes *IDH1*, *IDH2* and the patients with IDH mutations have a better prognosis (Yan et al. 2009; Houllier et al. 2010). We applied **Qombucha** separately on 896 IDH-WT GBM and 959 IDH-MUT patient samples, both of which are from TCGA. Because IDH-MUT cancers at any site are known to have disruptions in methylation (Nousmehr et al. 2010; Turcan et al. 2012) and there are also brain cancer-specific IDH-related methylation changes (Unruh et al. 2019), we choose to employ the completed reference panel *C_f_* obtained from the deconvolution of all 896 IDH-WT GBM samples, as described in Section ***Qombucha*** *recapitulates known GBM subtype characteristics*, to perform one-step QP-based simultaneous deconvolution of 896 IDH-WT GBM samples and 959 IDH-MUT samples. As can be observed from Figure 7A and 7B, IDH-WT GBM and IDH-MUT have different cell compositions and different favored developmental programs. Interestingly, IDH-WT GBM have higher overall presentation of the ASC-ODC-OPC-EC-VLMC progenitor and the “everything progenitor”, which are the two progenitors at the highest levels in the brain developmental tree. This suggests that IDH-WT GBM have greater stem-like characteristics than IDH-MUT, which potentially explains that IDH-WT GBM tends to have worse prognosis than IDH- MUT. Additionally, the *D_in_/D_ou_* scores presented in Figure 7C indicate that **Qombucha**-imputed cell fractions are able to distinguish IDH-WT GBM and IDH-MUT patients. Notably, IDH-MUT gliomas are known to originate from OPC (Rahme et al. 2023), and as can be observed in Figure 7A, our results mirror this prior finding.

**Figure 7:**
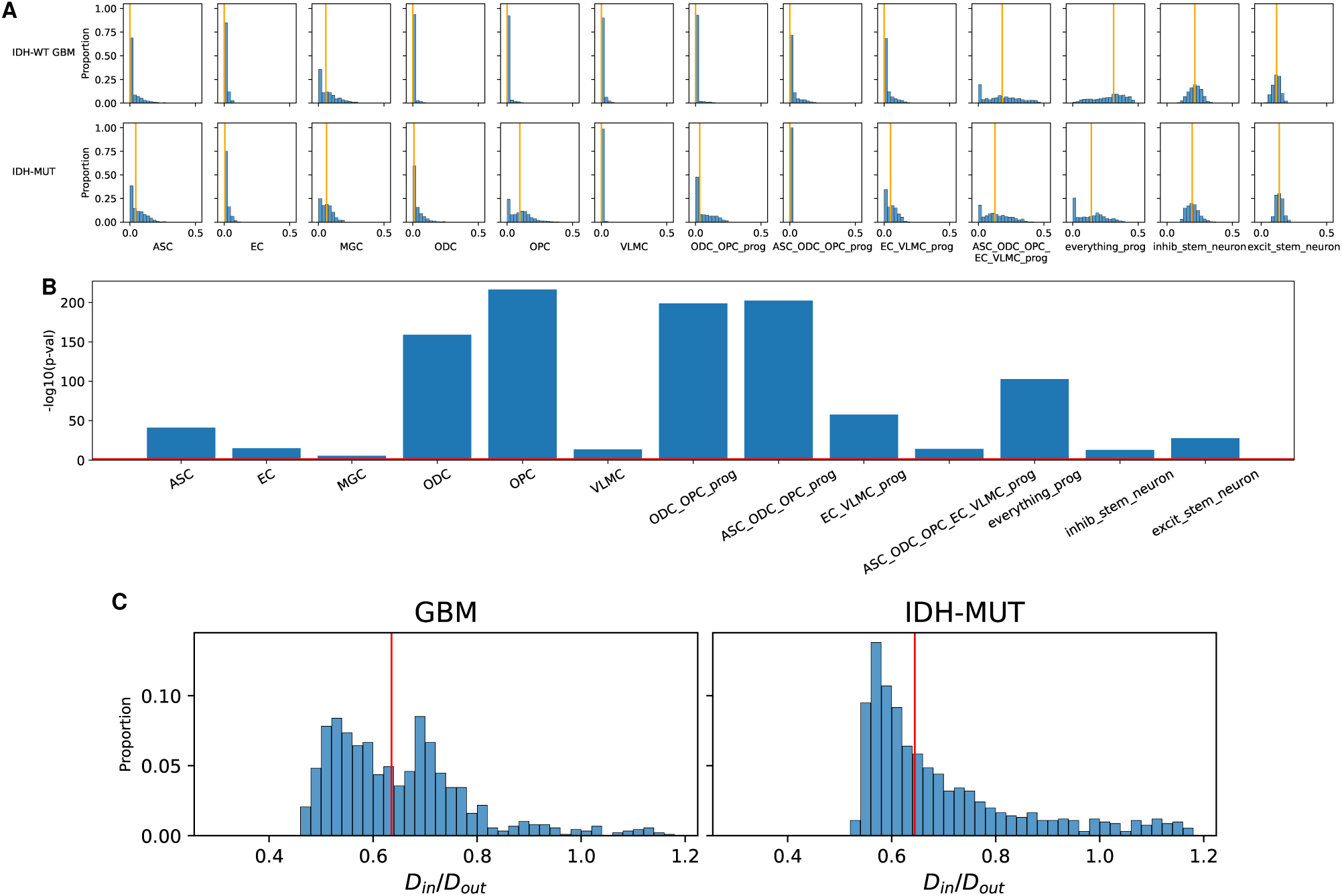
IDH-WT GBM and IDH-MUT gliomas have distinct cell type compositions. **(A)** Cell type compositions inferred by **Qombucha** in IDH-WT GBM and IDH-MUT. Orange vertical lines indicate the median inferred fraction for each cell type. Among the developed cell types, the primary difference between the IDH-WT GBM and IDH-MUT samples is in the fraction of OPC cells they harbor, as reported earlier (Rahme et al. 2023). **(B)** Bar plot showing *−* log_10_ p-values from KS tests on cell fractions inferred by **Qombucha** for IDH-WT GBM and IDH-MUT patients. The red horizontal line represents a p-value threshold of 0.05. Taller bars indicate more significant KS test results. **(C)** Tightness of IDH-WT GBM and IDH-MUT clusters evaluated using the *D_in_/D_out_* Score. The scores are calculated using the **Qombucha**-imputed cell fractions. The left panel shows the scores of the IDH-WI GBM patients, and the right shows that of the IDH-MUT patients.

### Qombucha-inferred progenitor methylation values are stably inherited

From the complete reference panel *C_ι_* output by **Qombucha**, we identified 39 features that are stably inherited (as defined in Section *Stable inheritance of methylation features*) along at least one of the five paths from the root of the brain development tree to a leaf with two or more edges. We performed the following simulation to show that this number (39) is higher than expected: we generated 1000 instances of randomly assigning methylation values (ranging between 0 and 1) to the unknown entries in the partially complete reference panel *C*, then evaluated how many methylation sites are stably inherited on at least one of the five paths. We repeated this process 5 times, and the fractions of the 1000 instances with 39 or more stably inherited sites on at least one path were 42/1000, 38/1000, 36/1000, 41/1000, 44/1000. See Figure 8A for one of the five experiments of the simulation; the proportion of replicates exceeding the threshold of 39 is the sum of the heights of the bars to the right of the vertical line. These results show that identifying 39 features stably inherited on at least one of the five paths is higher than expected, with an average empirical *P* = 0.0402. This demonstrates that on the brain development tree derived from the normal mature cell type methylation profiles by Tian et al. (2023), the progenitor cell types whose methylation profiles were identified on the TCGA GBM cohort by **Qombucha**, appear to stably inherit and transmit a high fraction (39 of the 98) of GBM-associated methylation features. Thus, from a methylation point of view, progression of the GBM cells is concordant with brain cell development trajectories.

**Figure 8:**
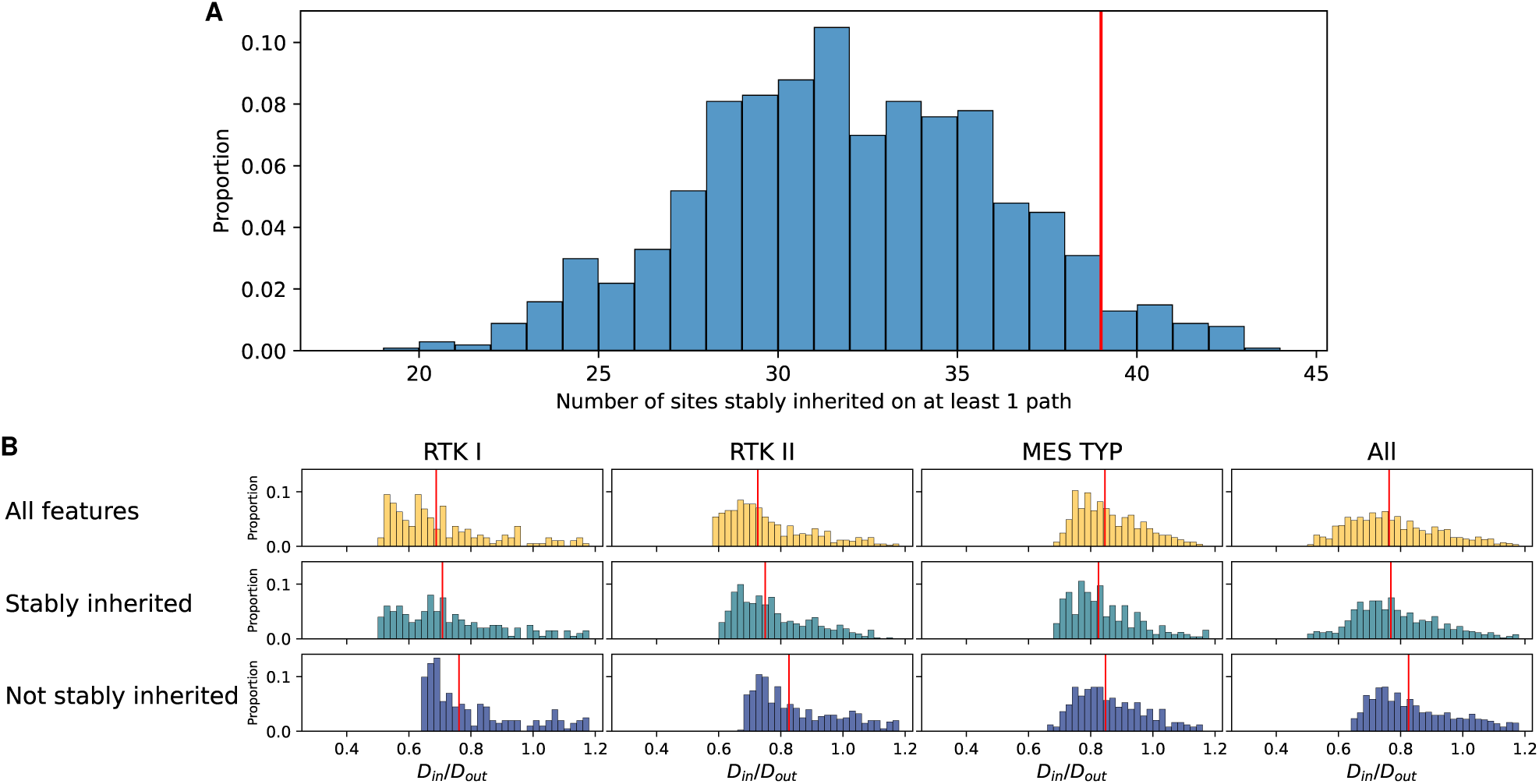
Qombucha inference identifies stably inherited methylation changes in brain development. **(A)** Random assignment of methylation value to evaluate the number of stably inherited sites obtained by chance. We generated 1000 instances of assigning random methylation values (between 0 and 1) to the unknown entries in the partially complete reference panel *C_p_*, then evaluated how many methylation sites are stably inherited on at least one of the five paths. The simulation was repeated 5 times, and one instance of the simulation is demonstrated in the distribution plot. The red vertical line marks 39, the number of stably inherited sites obtained from real data. **(B)** Tightness of GBM subtype clusters computed using *D_in_/D_out_*. Every row represents the tightness of known GBM subtype clusters computed using different information: all 98 features; 39 stably inherited features; 59 not stably inherited features. The red vertical lines indicate the median for each subtype in each experiment.

We then partitioned the **Qombucha**-inferred reference panel *C* into two disjoint subsets, one with 39 stably inherited features, the other with 59 not stably inherited features. Next, we inferred the cell fractions from the 896 TCGA GBM samples with one-step deconvolution using the two panels, respectively. Interestingly, the cell fractions inferred using only the stably inherited features demonstrate clustering tightness of the GBM subtypes comparable to when using all 98 features, whereas the cell fractions inferred using features not stably inherited show worse clustering of the subtypes, as can be seen in Figure 8B. These results demonstrate that stably inherited methylation changes in brain development dominate the distinctions among different GBM subtypes.

### Qombucha-inferred cell type proportions and association with overall survival

Among the 896 TCGA GBM samples (IDH-wildtype), overall survival (OS) was recorded for 299 samples. We hypothesized that having a larger proportion of defined cell types and a correspondingly smaller proportion of progenitor cell types would be be associated in our data with better overall survival because progenitor subtypes have been previously associated with tumor growth in other data Couturier et al. (2020). In Cox proportional hazards analysis, the sum of inferred proportions of the six defined (non-progenitor) cell types was almost significantly associated with OS at *P* = 0.05 and hazard ratio (HR) 0.2497 [0.0615, 1.003]. A related hypothesis is that the largest among the six inferred proportions of the six defined cell types would be associated with better overall survival and that was validated at *P* = 0.03 and HR = 0.1432 [0.02403,0.8536]. This suggests that the inferred cell type proportions have a statistical relationship to the clinically important measure of OS.

### Efficiency and robustness of Qombucha

We performed the following analyses to evaluate and validate the robustness and efficiency of **Qombucha**.

### Qombucha recovers simulated ground truth

We generated 5 random instances of the cell proportion matrix *P_sim_* and added a small Gaussiance noise to it, as described in Section *Simulations*. We then multiplied the randomly generated *P_sim_*with *C*_0_, the tree-informed intialization of reference panel, to obtain the simulated bulk mixture *B_sim_*. The goal is to evaluate whether **Qombucha** is able to recover the simulated ground truth cell proportions in *P_sim_*as well as the held out entries in *C*_0_. As can be observed in Figure 9A, the simulated cell fractions in *P_sim_* and **Qombucha**-imputed cell fractions, as well as the held-out entries of *C*_0_ and the **Qombucha**-imputed held-out entries, are both highly correlated. These results demonstrate the ability of **Qombucha** to solve for unknown entries in both *C* and *P* via the iterative process.

**Figure 9:**
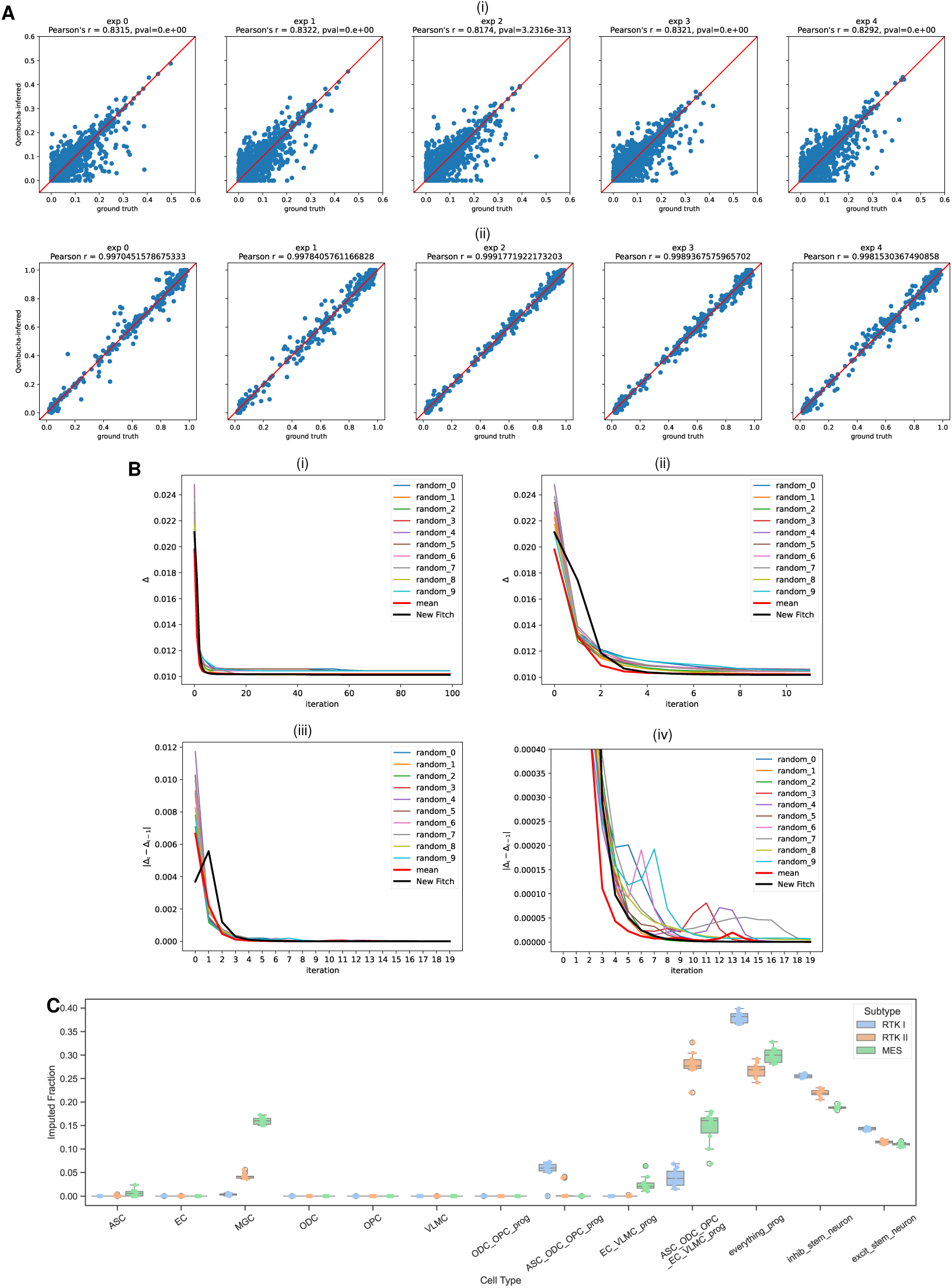
Qombucha is robust and efficient. **(A)** Correlation between values of simulated ground truth and **Qombucha**-inferred values. (i) Correlation of simulated cell fractions and **Qombucha**-inferred cell fractions. (ii) Correlation of held-out entries in *C*_0_ and **Qombucha**-inferred values of the held-out entries. **(B)** Informed starting point gives smaller objective value Δ and less volatile changes in Λ. Λ = Δ*_ι_ −* Δ*_ι_*_−1_. Random starting points are marked in thin lines. Starting point informed by averaging the mean of ancestors is marked in thick red lines. Starting point informed by the modified Fitch algorithm followed by the QP-based optimization is marked in thick black lines. (i) Change in Δ across iterations. (ii) Change in Δ across the first 10 iterations (zoomed-in version of (i)). (iii) Change in Λ across iterations. (iv) Change in Δ across the first 20 iterations (zoomed-in version of (iii)). **(C)** The distribution of median (across patients) inferred cell type fraction for each subtype across 10 random instances. Each instance involves randomly selecting 100 patient samples from each of the three subtypes to be used as the input matrix *B*.

### Qombucha executes numerous iterations rapidly and achieves quick convergence

The iterative update of matrices converges when the matrix reconstruction error is small and the change in objective function value between two consecutive iterations of **Qombucha** is below a user-defined threshold. The user-defined threshold parameters for the normalized matrix reconstruction error and the difference in objective value are denoted by *δ* and *λ*, respectively.

To evaluate the running time of **Qombucha**, we tested it with maximum iteration values of 100, 200, 300, 400, and 500 while setting the convergence tolerance parameters, *δ* and *λ*, to *−∞*. Using the bulk methylation matrix of 896 patients, processed as outlined in Section Pro*cessing bulk sample methylation data*, as well as the partial reference panel *C* and the initial reference panel *C*_0_ obtained as described in Section *Obtaining the partially complete reference panel C_p_ and initial reference panel C*_0_ *for non-neuronal cell types*, we observed that **Qombucha** completed 100 iterations in 23 minutes and 500 iterations in 114 minutes on a single CPU.

The choice of *δ* and *λ* may be critical in balancing runtime and solution accuracy. In our application of **Qombucha** to the TCGA GBM patient data, we observe that Δ decreases and stabilizes rapidly, as shown in Figure 9B(i) and 9B(iii), suggesting that running too many iterations may be unnecessary. After performing a parameter sweep, we noticed that setting *δ* to 0.0125 and *λ* to 10^−9^ offers a good trade-off between accuracy and runtime. With this setting, **Qombucha** converged within 86 iterations with the previously described bulk methylation matrix of 896 patients, partial *C* and initial *C*_0_ as input. For all **Qombucha** deconvolution of real patient data in this study, we set *max*_*iter*, the user-defined parameter specifying the maximum number of allowed **Qombucha** iterations to 100 while continuing to set *δ* to 0.0125 and *λ* to 10^−9^.

### Qombucha is robust to different starting points

We evaluated two alternative ways to fill in unknown entries in the initial value of the matrix *C_p_* to obtain *C*_0_ to test the robustness of iterative process of **Qombucha**, as descibred in Section *Alternative initialization of C*_0_: (i) random initialization, which is drawing values uniformly at random in the range [0, 1], and (ii) “mean of descendants” initialization, which is deterministically setting the methylation values of each row corresponding to a progenitor cell to be the average of its direct descendants in the tree of brain history development (Section *Reconstructing brain development history using single-cell methylation data* and Figure 3B for detail of the tree).

Random and mean starting points give comparable yet slightly higher objective value at convergence compared to the tree-informed QP-based method. The tree-informed QP-based initialization method is preferable because (i) it still has the best objective value at convergence and (ii) Λ = |Δ*_ι_ −* Δ*_ι_*_−1_| converges faster, as can be observed in Figure 9B.

### The brain development tree structure obtained via neighbor-joining outperforms alternative tree structures in GBM subtype clustering

In addition to the employed tree structure, which is obtained via neighbor-joining of the single-cell methylation data, we evaluated whether alternative plausible tree structures yield improved performance in GBM subtype clustering. Specifically, we enumerated all possible rooted binary trees, subject to the constraint that the clade ((OPC,ODC),ASC) is preserved in every tree. We apply this constraint because prior studies indicate that ASC and OPC share a common ancestor (Wang et al. 2025; Domingues et al. 2016), and the majority of differentiated OPCs turn into ODCs, suggesting a much closer developmental relationship between OPC and ODC (Nishiyama et al. 2014). Because the clade ((OPC,ODC),ASC) is fixed, the taxa {OPC,ODC,ASC} can be treated as one composite leaf. Thus, although there are originally six labeled leaves, each representing a developed cell type, the enumeration reduces to rooted binary trees on four leaves. The number of rooted binary trees with *n >* 1 labeled leaves is (2*n −* 3)!!(Felsenstein 1978); hence, the total number of trees consistent with this constraint is (2 *·* 4 *−* 3)!! = 5 *×* 3 *×* 1 = 15. The tree used in our study is among the 15 enumerated plausible trees.

We generated *C_p_* (see Section *Cell hierarchy-informed partial inference of progenitor cell type profiles*) and *C*_0_ (see Section *Cell hierarchy-informed initialization of unknown entries in the progenitor profiles*) for all 15 enumerated trees and solved for cell proportions using **Qombucha**’s iterative procedure with maximum iteration limit *max*_*iter* = 100. We then calculated the *D_in_/D_out_*scores (see Section *McClain–Rao score (D_in_/D_out_)*) using the cell proportions imputed from each tree structure: 0.7765, 0.7957, 0.7525, 0.7569, 0.7836, 0.7671, 0.7822, 0.7587, 0.7998, 0.7684, 0.7875, 0.7683, 0.7942, 0.7931, 0.8080. With the lowest *D_in_/D_out_* score 0.7525, the tree employed in our study outperforms all 14 other alternative tree structures in the clustering quality of GBM subtypes.

### Qombucha yields robust inferred cell compositions despite subsampling ***B***

From the bulk methylation matrix of 896 patients we processed as described in Section *Processing bulk sample methylation data*, we generated 10 instances of randomly subsampling 100 patient samples from each of the three subtypes and used each set of 300 patient samples as input. As shown in Figure 9C, the median inferred cell fractions of each subtype are largely consistent across the random instances. This result shows that **Qombucha**-imputed cell compositions are mostly robust to selection bias in input patient samples.

## DISCUSSION

In this study, we introduced **Qombucha**, a combinatorial optimization-based framework that leverages known brain developmental history to infer simultaneously methylation profiles of progenitor brain cells and cell type compositions from the methylation data of bulk GBM samples. We showed that **Qombucha** quickly produces optimal/near-optimal solutions and yields robust inference of cell type compositions despite added input noise or data subsampling.

Applying **Qombucha** to multiple GBM cohorts provided valuable biological insights, revealing that distinct brain developmental programs are associated with different GBM subtypes. We further demonstrated the generalizability of these findings by applying a TCGA-derived **Qombucha** reference panel to deconvolve an independent NCI GBM cohort and assigning GBM subtypes based on **Qombucha**-inferred cell-type compositions. A classifier trained on only 11 **Qombucha**-inferred cell-type fractions matched the performance of a state-of-the-art classifier based on the 10,000 most variable methylation sites. Furthermore, we demonstrated that **Qombucha**-inferred cell fractions per patient sample provide tighter clustering of GBM subtypes compared to other types of information, including the methylation values of 32k brain cancer subtype distinguishing methylation sites identified by Capper et al. (2018).

Together, these findings suggest that **Qombucha**-inferred cell-type compositions capture the core biological signals underlying GBM subtype differences while providing a substantially lower-dimensional and more interpretable representation of GBM tumor heterogeneity. Additional validation using single-cell data would strengthen these findings, though currently available datasets remain limited for this purpose. In particular, in one of the most comprehensive publicly available single-cell glioma datasets (Chaligne et al. 2021), the covered sites have very little overlap with the brain cell type–differentiating CpG sites analyzed in our study, limiting its utility for validation. Future single-cell methylation datasets with broader coverage of these CpG sites may enable more direct validation of our findings. In future work, we plan to explore how these developmental programs are linked to patient outcomes on other cohorts and to extend the application of **Qombucha** to other brain cancer subtypes.

## METHODS

### Simultaneous inference of cell type compositions and unknown methylation values in progenitor profiles

We formally define the problem of simultaneous inference of cell compositions and unknown methylation values in progenitor cells as a Quadratic Program (QP) below. The inputs to the QP are: (i) a matrix of bulk methylation data *B*, where each row corresponds to a vector of methylation *features* of a patient tumor sample, and (ii) a partially completed reference panel matrix *C_p_*informed by the fixed tree of brain development history with mature brain cells as leaves. Each feature is a set of methylation sites and the value of the feature in a sample is the sum of the methylation values at the sites in the feature.

In the framework introduced by Tian et al. (2023), a *feature* is the average methylation value between 0 and 1 among a set of individual methylation sites. To incorporate information from brain development, we first determine the methylation profiles of certain features in non-neuronal progenitors using the methylation profiles of mature brain cell types and the tree structure reflecting the history of brain development. The features of each progenitor whose methylation values can be determined are those that have consistent methylation values among all mature descendant cell types of the progenitor. The values of the remaining undetermined features in the progenitor profiles can be simultaneously inferred along with the cell composition using a quadratic programming (QP) formulation summarized in Figure 1. As described below, the aim of the QP is to deconvolve the input matrix *B* into the following two matrices: (i) *P* the cell type proportion matrix, and (ii) *C_f_* - the full reference panel matrix (i.e., the completed version of *C_p_*).

### Formulation of the Quadratic Program

The inputs to the QP-based deconvolution method include a bulk methylation profile matrix *B* of size *n × m*, where *n* is the number of patients and *m* the number of methylation features, and a partially filled reference panel matrix *C_p_* of size *r × m*, where *r* is the number of cell types. The reference panel matrix *C_p_*= *{c_k,j_}* consists of a stable methylation value per feature, for each mature and progenitor brain cell in the tree of brain development history. The elements of matrix *C_p_* are fully known for mature cell types, while those for progenitor cell types they may not be fully determined. The exact construction of *C_p_* is provided in the Section *Cell hierarchy-informed partial inference of progenitor cell type profiles*.

Let *P* = *{p_i,k_}* be the matrix of size *n × r* that stores the inferred fraction of each cell type for each patient, with *p_i,k_* being the inferred fraction of cell type *k* in patient sample *i*. In each patient sample *i*, the sum of cell fractions across all *r* cell types must be equal to 1. Let *U* be the set of indices of unknown entries in matrix *C_p_*. The value of every unknown entry *c_x,y_|*_(*x,y*)∈_*_U_* must be between 0 and 1. With || *X* || representing the Frobenius norm of matrix *X*, the simultaneous inference problem can be formulated as a QP as follows:

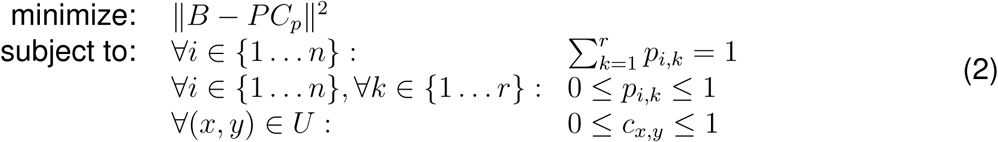

We use Gurobi (Gurobi Optimization, LLC 2024) (through its Python API) to solve the QP to simultaneously infer the estimated cell type compositions and the unknown entries in the partially complete progenitor profiles.

### Cell hierarchy-informed partial inference of progenitor cell type profiles

This section details the construction of the partial progenitor cell profiles from the methylation values of mature cells. Based on prior knowledge that GBM mostly originates from non-neuronal cells (Zong et al. 2015), we used the CpG sites and single-cell methylation reported by Tian et al. (2023) to construct a rooted binary neighbor-joining (Saitou and Nei 1987) tree for the 6 non-neuronal cell types reported in the same study, including astrocyte (ASC), oligodendrocyte (ODC), oligodendrocyte precursor (OPC), microglia (MGC), epithelial cell (EC), and vascular and leptomeningeal cell (VLMC) to reflect their development history (Figure 3B). In this tree, which we denote by *T*, the mature non-neuronal cell types are represented as leaves, and their presumptive progenitors as internal nodes. The differentiation of mature cells is represented as a branched process. The tree *T* we obtained has topologically differences from the cell hierarchy tree from Tian et al. (2023). In particular, it has a clade consisting of oligodendrocytes (ODC) and oligodendrocyte progenitors (OPC) that is biologically more plausible than the tree in Tian et al. (2023) in which ODC and OPC are apart. Even though OPC is the precursor of ODC, OPCs have distinct neural functions (Buchanan et al. 2023). In fact, many OPCs do not differentiate into ODCs (Kang et al. 2010). As a result, rather than inferring the developmental tree using only OPC (and not ODC) as a leaf and adding ODC as a child of OPC in a postprocessing step, we used both ODC and OPC as leaves of the developmental tree, and used the fact that OPC and ODC formed a clade within the developmental tree as a validation of our approach. A description of how *T* is constructed from the non-neuronal brain cell single-cell methylation profiles is in Section *Reconstructing brain development history using single-cell methylation data*, and how methylation feature values of the mature cell types, i.e., the leaves of *T* are determined can be found in Section *Obtaining the partially complete reference panel C_p_ and initial reference panel C*_0_ *for non-neuronal cell types*.

Next we infer the unknown methylation feature value for a progenitor, i.e. an internal node of *T*, so that it is “consistent” with the known values of the said feature across the progenitor’s descendants in *T* . Specifically, we identify for each internal node, the “range” of possible methylation values of the feature, from the node’s children, in a bottom-up manner (see Section *Obtaining the partially complete reference panel C_p_ and initial reference panel C*_0_ *for non-neuronal cell types* for details). We start by assigning to each feature of each leaf a range that includes only the known methylation value for that cell type. Then, for each internal node with children whose range of values for that feature has already been determined, we identify the smallest contiguous range that fully includes these two ranges; if this range is larger than a user defined threshold *δ*, we assign this range to the feature of the internal node otherwise, we calculate the median value of this range and assign to the feature of the internal node (the range that only includes) this median value. This algorithm is called “modified Fitch” because it is inspired by the algorithm in (Fitch 1971).

The pseudocode for this process is given in the “bottom-up phase” of Algorithm 2 (until Line 22). When the process terminates at the root of the tree, if any given feature of an internal node is assigned a single value, this value will be fixed. If instead, the feature is assigned a range of values for this internal node, a specific value from this range will be determined by **Qombucha**. The input to **Qombucha** is thus these partially determined feature values, represented as reference panel *C_p_* of size *r × m*, where *r* is the number of cell types, *m* is the number of methylation features, and thus row *c_k_* is the methylation profile of cell type *k*. *C_p_* has a fixed value for each feature of a mature cell type, and some of the features of each progenitor; it also specifies the range of values that an unknown feature of a progenitor can take.

### Qombucha: Solving the QP Iteratively

A direct simultaneous inference of the cell composition matrix *P* and unknown entries in *C_p_* in the quadratic program (2) requires substantial time and computing resources to find the optimal solution. To improve computational tractability, we introduce **Qombucha**, a block coordinate descent-like approach based on iteratively updating cell type composition and unknown entries in progenitor profiles, as summarized in Figure 2. An open-source implementation of **Qombucha** is available at: https://github.com/algo-cancer/Qombucha.

### Iterative updating of *P* and unknown entries in *C*

Instead of trying to solve unknown entries in *C_p_* and the cell composition matrix *P* simultaneously, **Qombucha** initializes the unknown entries guided by *T* to obtain *C*_0_ and then uses *C*_0_ as a starting point in an iterative procedure to alternate between updating *P* and the unknown entries in *C_p_* until convergence. The construction of *C*_0_ is described in Section *Cell hierarchy-informed initialization of unknown entries in the progenitor profiles*. In each iteration *ι*, we first solve a restricted version of the QP described in Section *Formulation of the Quadratic Program* for *P_ι_* with *B and* fully known *C_ι_*_−1_ as inputs:

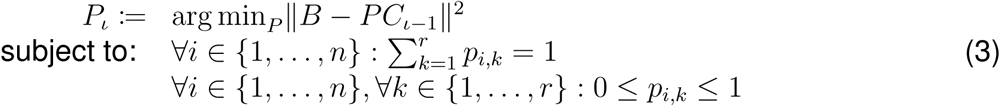

Then, we solve another restricted version of the QP described in Section *Formulation of the Quadratic Program* for the unknown entries in *C_p_*(whose indices form the set *U*) to obtain *C_ι_* with *B* and *P_ι_* (computed above) as inputs:

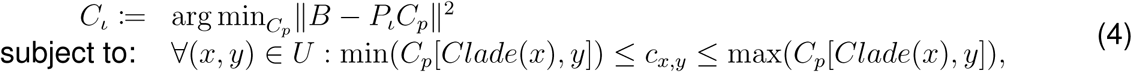

where *Clade*(*v*) is the set of all leaf labels in the subtree of *T* rooted at vertex *v* and *C_p_*[*Clade*(*v*)*, j*] *{C_p_*[*i, j*] *| ∀i ∈ Clade*(*v*)*}*. The unknown entries of each progenitor cell are constrained by the methylation values of their descendants for the following reasons. As DNA methylation is binary at the allele level, the measured methylation level of a CpG site for a particular cell type reflects the fraction of methylated alleles across a cell population of that type. A progenitor population that gives rise to two or more differentiated cell types can be viewed as a mixture of subpopulations already biased toward each fate. As the methylation features identified by Tian et al. (2023) are composed of a set of sites which are lineage informative, for any given feature, a progenitor cell type’s average methylation level is expected to be between the methylation levels of its descendants.

**Qombucha** outputs *P_ι∗_* and *C_ι∗_* provided it “converges” at some iteration *ι*^∗^. The algorithm is said to have converged when any of the following conditions is satisfied: (i) the bulk methylation matrix reconstruction error is “small” and the change in objective function value between successive iterations is “small”, or (ii) the algorithm has run long enough. Specifically, in each iteration *ι*, **Qombucha** minimizes the squared error between the observed bulk methylation data matrix *B* and the reconstructed matrix *B_ι_* = *P_ι_C_ι_*. At the end of iteration *ι*, the normalized squared difference, Δ*_ι_*, is calculated between *B* and *B_ι_* as:

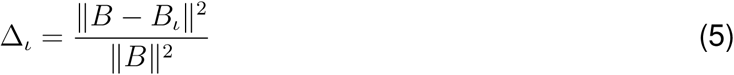

**Qombucha** is said to have converged at iteration *ι* if Δ*_ι_* falls below a user-defined *δ* and Λ = Δ*_ι_ −* Δ*_ι_*_−1_ is smaller than a user-defined *λ*; this is based on the expectation that another iteration would be unlikely to reduce the error substantially. Additionally, **Qombucha** has a user-defined maximum iteration limit *max*_*iter*, which is set to bound the running time. The pseudocode for the iterative procedure is given in Algorithm 1.

Algorithm 1 Iterative optimization for P and unknown entries in C

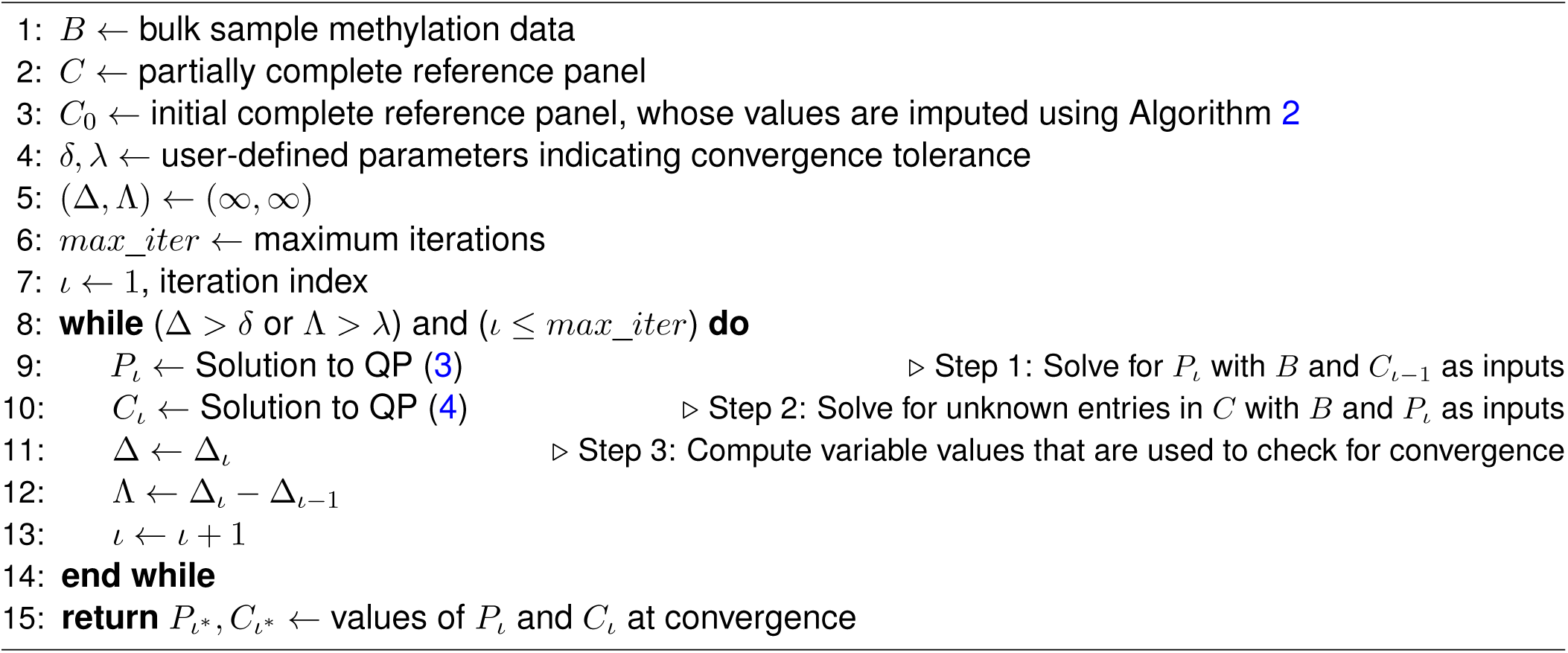

### Cell hierarchy-informed initialization of unknown entries in the progenitor profiles

The partially complete reference matrix *C_p_* is obtained as described in Section *Cell hierarchy- informed partial inference of progenitor cell type profiles*. We utilize the brain cell development tree *T* to inform the initialization of “unknown entries” in *C_p_* to obtain the initial complete reference matrix *C*_0_, which serves as the starting point in the iterative procedure of **Qombucha**. The *unknown entries* in matrix *C_p_* are those entries corresponding to the progenitor cells whose feature values are left as ranges at the end of the bottom-up phase of Algorithm 2. If, for any methylation feature *y* and progenitor cell type *x*, the methylation range of each unknown entry in *C_p_* is given by [*m_low_*[*x, y*]*, m_high_*[*x, y*]], then the optimal initial methylation values are simultaneously determined by minimizing the sum of squared methylation changes between parent-child pairs across the entire tree, using the following quadratic program:

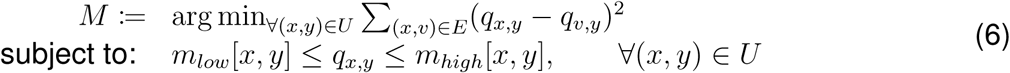

where *q_x,y_* represents the methylation level at the parent node for feature *y* and *q_v,y_* represents the methylation level at the child node for the same feature. The complete pseudocode for building *C_p_*and obtaining *C*_0_, is given in Algorithm 2.

Algorithm 2 Initializing unknown entries in *C_p_* for a given methylation feature *s*

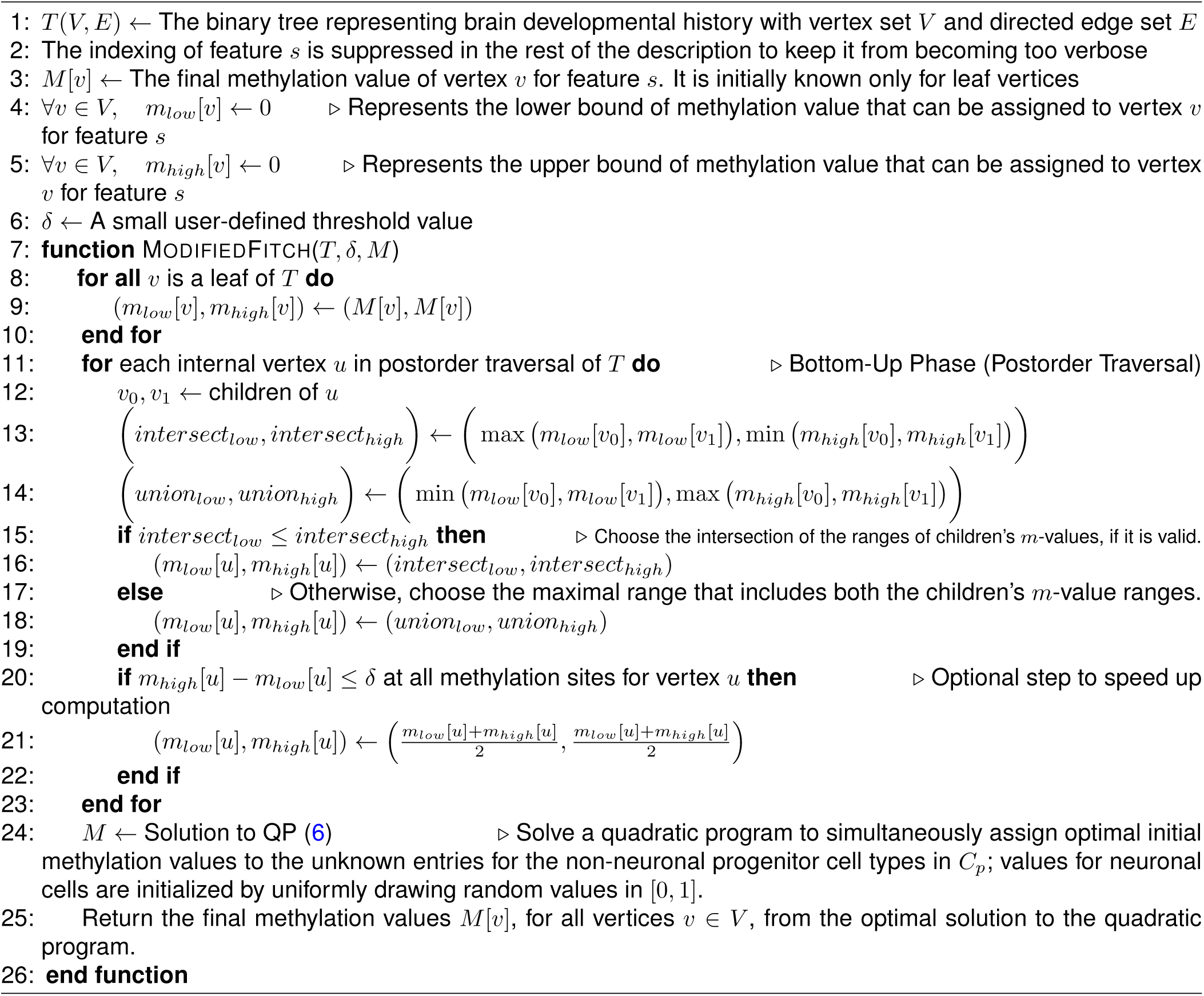

### Evaluating tightness of the known GBM subtype clusters

To evaluate how well different types of information (i.e., **Qombucha**-imputed cell fractions, the methylation values of Capper et al. (2018)-reported 32k sites etc.) recapitulate the known GBM subtype clusters, we use the McClain–Rao score (McClain and Rao 1975), i.e., the ratio between the average in-class distance and the average out-of-class distance (*D_in_/D_out_*) as well as the silhouette score (Rousseeuw 1987) to assess the tightness of clusters.

### McClain–Rao score (*D_in_/D_out_*)

This score corresponds to the ratio between the mean in-class distance and mean out-of-class distance for each patient to evaluate the tightness of GBM subtype clusters. Let *v_i_* be the feature vector representing patient sample *i*, and assume that sample *i* belongs to class *θ*. Let *α* denote the number of samples in class *θ*, and *β* denote the number of samples outside of class *θ*. The mean *L*^1^ distance between sample *i* and other samples within the same class *θ* is defined as:

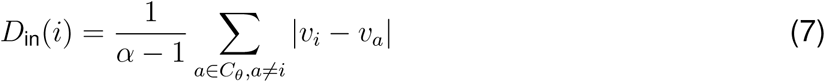

where *C_θ_*represents the set of samples belonging to class *θ*, and *a* indexes over all samples in *C_θ_* excluding sample *i*. The mean *L*^1^ distance between sample *i* and samples outside of class *θ* is defined as:

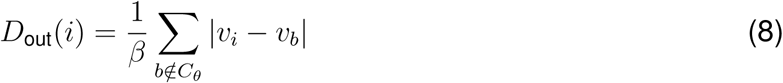

where *b* indexes over all samples outside class *θ*. The clustering goodness for sample *i* is then defined as the ratio of the in-class distance to the out-of-class distance:

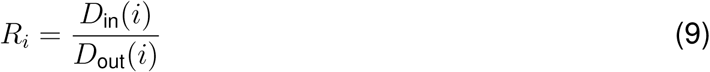

*R_i_* is the ratio that indicates the tightness of the clusters: a value close to 0 indicates tighter clustering (i.e., the mean in-class distance is much smaller than the mean out-of-class distance), while a value close to 1 indicates poor clustering and a value larger than 1 indicates that sample *i* has a closer affinity with samples outside of its assigned class *θ* than the samples within class *θ*.

### The silhouette score

A silhouette value *s*(*i*), as defined by Rousseeuw (1987), can be computed for each item *i* in any given cluster as follows: Let *a*(*i*) denote the average dissimilarity of the item to all other items in its own cluster *C_I_*, and *b*(*i*) denote the average dissimilarity of item *i* to all items in its closest neighboring cluster *C_J_* . That is,

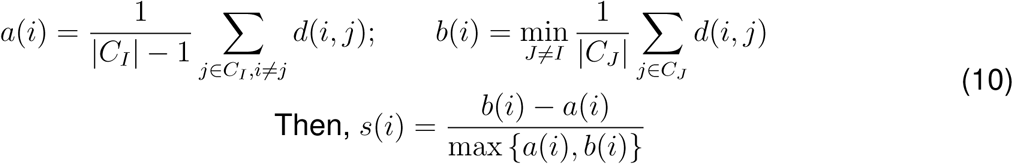

Here, *d*(*i, j*) could be any distance metric. We use *L*^1^ norm in our evaluation (our observations hold true for *L*^2^ norm as well). By definition, *s*(*i*) lies in [*−*1, 1]; the closer the value is to 1, the better clustered item *i* is. In contrast, if *s*(*i*) is closer to *−*1, it would be more appropriate for item *i* to be taken out from its current cluster and assigned its closest neighboring cluster. We compute *s*(*i*) value for each patient *i* using the corresponding row vectors from *P_ι∗_* as features.

### Input data preparation

#### Reconstructing brain development history using single-cell methylation data

We randomly selected 30 cells per each non-neuronal cell type from the Tian et al. (2023) dataset. The sampling ensures that each cell type has equal representation in the cohort to avoid biasing the tree. As explained above, the value of a feature in a sample is the average methylation value for sites used in that feature group for each feature group reported in Tian et al. (2023). The pairwise distances of the cells are calculated based on the values of the 800 feature groups. The tree representing the branched process underlying brain development history is then constructed using neighbor-joining (Saitou and Nei 1987) based on the pairwise distances. The resulting tree structure is shown in Figure S1. As can be observed from the tree, the cells of the same type cluster together with only a few exceptions. We then infer the brain developmental history by contracting the clades of the same cell types. Additionally, we removed the cells denoted “PC” from the brain development history as this cell type is inconsistently defined in the Tian et al. (2023) manuscript and the provided data labels. In particular, in the supplement of Tian et al. (2023) “PC” is defined as “perivascular cells”, but in the metadata associated with the single-cell methylation profiles, “PC” is defined as generic cells (cell ontology ID CL:000000). The resulting brain development history is depicted in Figure 3B.

#### Processing bulk sample methylation data

We analayzed two datasets referred to as TCGA and NIH samples, respectively. For the TCGA samples, we started with bulk methylation data for 1,160 samples sequenced using the Infinium HumanMethylation450 BeadChip (450k array) from the TCGA database, which were deemed to be sufficiently complete for tumor subtype classification analysis by the NCI/CCR Laboratory of Pathology. The samples were classified as RTK I, RTK II, or MES TYP subtypes using the Capper et al. (2018) brain and CNS tumor classifier. To eliminate potential biases stemming from tumor impurity and subtype classification errors, we removed samples with lower than 0.5 tumor purity and subtype assignment scores lower than 1, resulting in 896 patient samples passing quality control. Among the 896 samples, there are 216 RTK I samples, 431 RTK II samples, and 249 MES TYP samples. The 450k methylation array data cover 485,512 sites, among which 2,484 sites overlap with the 156k CpG sites reported by Tian et al. (2023). The 2,484 sites were then grouped according to the features reported in Tian et al. (2023), resulting in 98 brain development-associated features with at least one site among the original 800 features reported in Tian et al. (2023). The other 702 features reported in Tian et al. (2023) are excluded, as none of the 485k methylation array sites overlap with the brain cell type-differentiating CpG sites in these feature groups. We similarly processed 1,056 IDH-mut glioma 450k array samples to obtain 959 samples passing quality control.

The NIH GBM dataset comprises 556 patient samples profiled using the Illumina Human Methylation EPIC 850K DNA methylation array (EPIC 850k array). Among the 556 samples, there are 112 RTK I samples, 186 RTK II samples, and 258 MES TYP samples. Genomic DNA extracted from FFPE tumor tissue underwent bisulfite conversion and FFPE DNA restoration before analysis on the Illumina Infinium Methylation array, following the protocol described previously (Chung et al. 2024). Samples included in this analysis were collected through May 13, 2025. Raw intensity values from the EPIC 850K DNA arrays were read into data objects using the minfi R library (Aryee et al. 2014). Preprocessing was implemented as described in Capper et al Capper et al. (2018). Specifically, probe intensity values were normalized using the preprocessIllumina function. Probes were filtered to exclude those with ambiguous mappings to the reference genome, those representing non-CpG sites, those representing single nucleotide polymorphisms, those located on the sex chromosomes, and those shown to produce misleading results. As a result, 702,068 sites, out of 853,307 sites covered in the EPIC 850k array, passed quality-control filtering. To enable direct comparison with the TCGA dataset, we intersected the quality-controlled EPIC v1.0 sites with the 2,484 sites shared between the Illumina 450K methylation array and the 156k CpG sites reported by Tian et al. (2023). This overlap yielded 1,932 CpG sites, which were then grouped according to the feature definitions described in Tian et al. (2023). Among the original 800 feature groups, 93 brain development-associated features containing at least one CpG site were retained in the NIH dataset; these 93 features represent a subset of the 98 features retained in the TCGA dataset. The remaining 707 feature groups reported in Tian et al. (2023) were excluded, because none of the 1,932 overlapping CpG sites corresponded to the brain cell type-differentiating CpG sites within those feature groups.

The NIH samples were obtained from patients undergoing treatment for which they gave informed consent at hospitals with which the NCI Laboratory of Pathology has collaboration agreements to receive patient samples for secondary analysis. From the NIH Institutional Review Board (IRB), we (KDA) obtained a waiver for additional informed consent for secondary use of the samples in research, as is done in this study.

#### Obtaining the partially complete reference panel *C_p_* and initial reference panel *C*_0_ for nonneuronal cell types

We combined the single-cell methylation profiles of 30 cells randomly sampled from each non-neuronal cell type (ASC, ODC, OPC, MGC, EC, and VLMC) and computed the average methylation value for sites in each of the 98 methylation feature groups selected by Tian et al. (2023). This resulted in a reference panel of size 6 *×* 98 for mature non-neuronal cells. Then, using a threshold of *δ* = 0.01 and cell hierarchy-informed inference of progenitor cell profiles as described in Section *Cell hierarchy-informed partial inference of progenitor cell type profiles*, we obtained the partially complete profile of non-neuronal progenitors. Appending the profiles of mature non-neuronal cells and the partially inferred progenitors, we obtained the partially complete reference panel *C_p_* of size 11 *×* 98 (see Figure S2A). *C_p_* contains 327 unknown entries. To obtain the non-neuronal entries of initial reference panel *C*_0_, which serves as the starting point in the iterative procedure of **Qombucha**, we initialized the 327 unknown entries in *C* using the method described in Section *Cell hierarchy-informed initialization of unknown entries in the progenitor profiles* guided by the brain development history *T* constructed as described in Section *Re- constructing brain development history using single-cell methylation data* (see Figure S2B). To complete *C_p_* and *C*_0_ we added two neuronal cell types as described in Section *Expanding C_p_ and C*_0_ *with neuronal cell types*.

#### Expanding *C_p_* and *C_0_* with neuronal cell types

While GBM is a cancer of non-neuronal cells, GBM samples may include impurities consisting of neuronal cells, as non-neuronal cells like astrocytes (ASC) and oligodendrocytes (ODC) can be in close proximity of neuronal cells (Auld and Robitaille 2003). Therefore, we also add to *C_p_* and *C*_0_ the partially complete excitatory and inhibitory neuronal cell consensus methylation profiles. In the Tian et al. (2023) dataset, 15 cell types are in the excitatory neuronal cell category, and 18 in the inhibitory neuronal cell category. Similar to the processing of non-neuronal cells, we randomly sampled and combined the single-cell methylation profiles of 30 cells of each neuronal cell types. Then, for the excitatory and inhibitory neuronal cell categories, respectively, we computed the partially complete consensus profiles by averaging values of methylation features consistent across cell types within the same category (standard deviation *≤* 0.01) and leave inconsistent methylation features (19 in the excitatory category and 52 in the inhibitory category) empty. After including the excitatory and inhibitory neuronal cell consensus profiles, *C_p_* has size 13 *×* 98. The empty entries of the neuronal consensus profiles are then initialized by randomly drawing values in the range [0, 1] and the resulting two rows are included in *C*_0_, making it also of size 13 *×* 98.

### Experiments for robustness and benchmarking

#### Simulations

We generated *P_sim_* by randomly drawing 100 samples, each with 13 dimensions, the number of cell types included in this paper, from a symmetric Dirichlet distribution with concentration parameter *α* = 1 for all 13 dimensions. The randomly generated *P_sim_* is then multiplied with *C*_0_, which was generated as described in Sections *Obtaining the partially complete reference panel C_p_ and initial reference panel C*_0_ *for non-neuronal cell types* and *Expanding C_p_ and C*_0_ *with neuronal cell types*, followed by introducing Gaussian noise (*µ* = 0*, σ* = 0.01) to obtain *B_sim_*. The entries in *B_sim_* are corrected to 0 if *<* 0 and 1 if *>* 1.

#### Alternative initialization of *C*_0_

To test the usefulness of the algorithm-guided initialization of matrix *C*, i.e., the matrix *C*_0_ obtained through Algorithm 2, we compare the quality of solutions to **Qombucha** instances against two other modes of initialization: (i) random initialization the unknown entries of the matrix *C*_0_ are filled by drawing real numbers uniformly within range [0, 1]; (ii) “mean of descendants” initialization the unknown entries of *C*_0_ are filled by setting the methylation values of each internal node in the tree *T* to be the mean of its direct descendants’ values.

#### Subsampling patient samples

From the bulk methylation matrix of 896 GBM patients we processed as described in Section *Processing bulk sample methylation data*, we generated 10 patient subsets, each by randomly subsampling 100 patient samples from each of the three subtypes (RTK I, RTK II, and MES TYP). Each subset of 300 patient samples is then used as input to **Qombucha** to test its robustness.

#### Benchmarking quality of clustering

We benchmarked the quality of clustering of **Qombucha**-inferred cell fractions by computing McCain-Rao scores (*D_in_/D_out_*) and silhouette scores (described in Section *Evaluating tightness of the known GBM subtype clusters*) using the following information: (i) the methylation values of 2484 sites randomly sampled from the 450k methylation array, (ii) the methylation values of Capper et al. (2018) 32k sites, (iii) the methylation values of 98 features containing cell type- differentiating sites included in the 450k array (see Section *Input data preparation* for a detailed explanation), (iv) QP-inferred non-neuronal mature brain cell fractions, excluding progenitor profiles, (v) MethylCIBERSORT(Chakravarthy et al. 2018)-inferred non-neuronal mature brain cell fractions, excluding progenitor profiles, and (vi) **Qombucha**-inferred non-neuronal mature and progenitor cell fractions with alternative initialization of *C*_0_ as described in Section *Alternative initialization of C*_0_.

#### Stable inheritance of methylation features

We define a methylation feature to be *stably inherited* in a given path along the brain development tree, if the overall standard deviation of the methylation values on the path is larger than or equal to 0.01, and the methylation value is strictly monotonic along the path.

#### Wilcoxon rank-sum test for evaluating similarity of cell proportions across cohorts

To assess the similarity of inferred cell-type proportions between the TCGA GBM and NCI GBM cohorts, we compared the distributions of cell-type proportion rankings across patient cohorts using Wilcoxon rank-sum tests. Because the two cohorts contained different numbers of patients within each GBM subtype, we randomly sampled 100 patients per subtype from each cohort. This sampling procedure was repeated 100 times.

For each instance of random sampling, inferred proportions for each cell type were ranked across all sampled patients within a cohort, irrespective of subtype. Subsequently, within each GBM subtype and for each cell type, Wilcoxon rank-sum tests were performed to compare the ranked cell-type proportions between the two cohorts. Across 13 cell types and 3 GBM subtypes, a total of 39 comparisons were conducted. Statistical significance was therefore determined using a Bonferroni-corrected threshold of 0.05/39. All statistical analyses were performed using the scipy.stats.wilcoxon function in SciPyVirtanen et al. (2020) with the parameter setting zero_method=“zsplit”.

#### Survival Analysis

Analysis of the possible relationship between overall survival (OS) and **Qombucha**-inferred cell type proportions was done with the Cox proportional hazards method as implemented in the coxph function of the R library survival Therneau and Grambsch (2000).

#### Classification of GBM subtypes using Qombucha-inferred cell fractions

To classify GBM subtypes based on inferred tumor composition, we developed an ensemble classification framework using **Qombucha**-imputed cell type fractions as input features. For each sample, the feature vector consisted of the inferred proportions of cell types obtained from Qombucha. Prior to model training, all features were scaled to the range of [0, 1] using the MinMaxScaler function from scikit-learn. We evaluated four classification algorithms: k-nearest neighbors (KNN), logistic regression (LR), random forest (RF), and support vector machine (SVM). Each model outputs class probabilities corresponding to GBM subtypes (RTK I, RTK II, MES TYP). We averaged the predicted class probabilities across all four models. A sample was assigned to the class with the highest average probability and that highest average probability was used as a real number of ROC-AUC analysis.

Classification performance on the TCGA cohort was assessed using a 5 × 5 nested cross-validation framework following a protocol as previously described in Hoang et al. (2024). Briefly, the TCGA cohort was partitioned into five outer folds, with each outer fold serving once as an independent test set (20% of samples), while the remaining four outer folds (80% of samples) were used for model training. Within each outer training set, a second five-fold cross-validation was performed, in which each inner fold was sequentially held out as a validation set for model selection and hyperparameter optimization. This procedure generated five models for each outer fold, resulting in a total of 25 trained models. For external validation on the independent NCI cohort, predictions from all 25 models were averaged to generate the final subtype prediction, without additional training or parameter optimization. Model generalizability was evaluated on an independent NCI cohort. Model performance was assessed using overall accuracy and the area under the receiver operating characteristic curve (AUC), which was computed using a one-vs-rest strategy and macro-averaged across classes. Confusion matrices were used to further evaluate classification performance across subtypes.

#### Classification of GBM subtypes using highly variable methylation CpGs

To classify GBM subtypes using DNA methylation profiles, we selected the 10,000 most variable CpG sites across samples in the TCGA GBM cohort based on variance in beta values. These highly variable CpGs were used as input features for subtype classification.

A support vector machine (SVM) classifier was trained using the TCGA GBM cohort with subtype labels corresponding to RTK I, RTK II, and MES TYP. To assess model generalizability, the trained SVM classifier was independently evaluated on the NCI cohort. Classification performance was quantified using overall accuracy and the area under the receiver operating characteristic curve (AUC). Multi-class AUC values were computed using a one-vs-rest approach and macro-averaged across subtypes. Confusion matrices were additionally generated to evaluate subtype-specific classification performance.

## DATA ACCESS

### Lead contact

Requests for further information and resources should be directed to and will be fulfilled by the corresponding author, S.Cenk Sahinalp.

## Data availability

The processeed TCGA data is available upon request. The NCI-generated data cannot be shared because the human study subjects on whom methylation data were collected at NIH did not consent to data sharing.

## ACKNOWLEDGMENTS

This research was supported in part by the Intramural Research Program of the National Institutes of Health (NIH). The contributions of the NIH authors were made as part of their official duties as NIH federal employees, are in compliance with agency policy requirements, and are considered Works of the United States Government. However, the findings and conclusions presented in this paper are those of the authors and do not necessarily reflect the views of the NIH or the U.S. Department of Health and Human Services. This work utilized the computational resources of the NIH HPC Biowulf cluster (https://hpc.nih.gov).

## Author Contributions

Conceptualization, XCL, AAS, and SCS; methodology, XCL, HL, AH, AAS, and SCS; investigation, all authors; writing–original draft, XCL, HL, AH, AAS, and, SCS; writing–review & editing, all authors; funding acquisition, SCS; resources, KDA, ER, SCS; supervision, SMM, KDA, ER, AAS, SCS.

